# Induction of a CD8 T cell intrinsic DNA damage and repair response is associated with clinical response to PD-1 blockade in uterine cancer

**DOI:** 10.1101/2022.04.16.488552

**Authors:** Yuki Muroyama, Sasikanth Manne, Nils Wellhausen, Derek A. Oldridge, Allison R. Greenplate, Lakshmi Chilukuri, Divij Mathew, Caiyue Xu, Ramin S. Herati, Shelley L. Berger, Alexander C. Huang, Carl H. June, Dmitriy Zamarin, Claire F. Friedman, E. John Wherry

## Abstract

Despite the success of immune checkpoint blockade (ICB), many patients fail to achieve durable clinical benefit, and the underlying immunological mechanisms remain poorly understood. Here, we investigated immune reinvigoration by ICB in advanced or recurrent hypermutated or microsatellite instability-high, mismatch repair deficient (MSI-H/MMRd) uterine cancer patients treated with anti-PD-1 (nivolumab). CD8 T cells underwent rapid pharmacodynamic proliferation 2-4 weeks after initiating PD-1 blockade. This immunological response, however, did not correlate with clinical response. We hypothesized that the T cell-intrinsic response to proliferative and genotoxic stress might contribute to the disparity between immunological and clinical response. We developed a high-dimensional single cell cytometric platform to simultaneously analyze T cell differentiation with changes in DNA damage and repair (DDR) pathways. This DDR-Immune platform revealed T cell subset-specific patterns of DDR, and distinct DDR pathways induced by different types of DNA damage. Applying this platform to MSI-H/MMRd or hypermutated uterine cancer patients revealed a signature of DDR exemplified by rapid increase in phosphorylated-ATM (pATM) intrinsic to CD8 T cells proliferating in response to PD-1 blockade that distinguished clinical responders and non-responders. ATM regulated transcriptional circuits in T cells were associated with better clinical response to PD-1 blockade. These findings highlight a previously unrecognized role for CD8 T cell-intrinsic DDR as a potential determinant of immune fitness and clinical outcome of PD-1 blockade therapy.

**STATEMENT OF SIGNIFICANCE:** Using high-dimensional immune and DNA damage response (DDR) profiling in T cell subsets we identified distinct signatures in anti-PD-1 treated MSI-H/hypermutated uterine cancer patients associated with clinical response versus non-response. Thus, T cell-intrinsic DDR is a potential determinant of immune responsiveness and clinical outcome to PD-1 blockade therapy in cancer.

## INTRODUCTION

Despite the success of immune checkpoint blockade (ICB), many patients still fail to achieve durable clinical benefit and the underlying immunological determinants of clinical response versus non-response remain poorly understood. Although many components of the immune system likely contribute to successful responses to ICB in cancer, accumulating evidence implicates CD8 T cells, including exhausted CD8 T cells (T_EX_), in the immunological effects of PD-1 pathway blockade(1–6). For some cancers such as melanoma, lung cancer, and basal cell carcinoma, T_EX_ are reinvigorated by PD-1 pathway blockade and these reinvigorated T_EX_ can contribute to disease control (7–11). This immunological response to ICB can be monitored in peripheral blood and is defined by the rapid and robust induction of proliferation in the pool of circulating T_EX_ and possibly other types of T cells. This response involves pre-existing T_EX_ specific for tumor antigens, including neoantigens, but may also result in new T cell priming and *de novo* T cell responses in some patients (8,12,13). Nevertheless, not all patients with this “immunological response” to PD-1 blockade experience an effective clinical response (7–11). The underlying mechanisms for suboptimal T cell responses to anti-PD-1 therapy remain incompletely understood.

Upon proper activation, antigen-specific T cells undergo rapid proliferation (14–16) with cell cycle times as short as 2-4 hours (17). This rapid cell cycle occurs in the setting of cell intrinsic production of reactive oxygen species from robust mitochondrial metabolism and often environmental free radicals due to inflammation. These environmental conditions and proliferative stress have the potential to cause genomic damage to T cells responding to simulation without allowing sufficient time to repair subsequent DNA damage, leading to exit from cell cycle, senescence, and/or loss of proliferative capacity (18, 19). To maintain genome integrity, rapidly proliferating T cells must be equipped with cell-intrinsic DNA damage and repair (DDR) mechanisms. We recently described a role for such cell intrinsic DDR response capacity in a subset of CD8 T cells in mice early after initial T cell activation where this DDR capacity was necessary for optimal CD8 T cell memory formation(20). Moreover, these studies identified a link between T cell intrinsic DDR and inhibitory receptor pathways including PD-1 (20). Whether such a role for DDR also exists for human CD8 T cells and how different subsets of human CD8 T cells may differ in DDR is unknown. In addition, it is unclear how such cell intrinsic DDR in immune cells may impact response to immunotherapy in cancer, where T cell proliferation is a requisite feature of immunological response to immunotherapies such as ICB. This question may be particularly relevant in patients who have been pre-treated with DNA damage-inducing therapies, such as chemotherapy, targeted DNA repair inhibitors (e.g. PARP inhibitors), radiotherapy, or in whom advanced age is associated with altered T cell differentiation and senescence. Moreover, features of some cancers themselves, such as the overall burden of mutations or antigens driving immune stimulation could also impact the requirements for T cell-intrinsic genomic surveillance.

To begin to investigate how T cell-intrinsic DDR may relate to the immunological and clinical response to PD-1 blockade in cancer immunotherapy, we examined a cohort of microsatellite instability-high, mismatch repair deficient (MSI-H/MMRd) or hypermutated (≧ 20 mutations/megabase (Mb)) uterine cancer patients treated with nivolumab. In general, patients with MSI-H/MMRd or hypermutated tumors have high TMB and respond better to PD-1 blockade (21–23), likely due to the generation of neoantigens that can be recognized by T cells (23–28). Despite this association of TMB with clinical response to anti-PD-1 therapy, however, many patients still fail to achieve durable clinical responses, suggesting additional immune dysregulation (23). Because this uterine cancer cohort had high TMB, and most patients mounted a proliferative response to PD-1 blockade this cohort provided an ideal opportunity to explore the gap between immunological and clinical response to PD-1 blockade and to test the impact of T cell-intrinsic DDR. Our immune profiling of this MSI-H/MMRd or hypermutated uterine cancer cohort revealed a clear immune-pharmacodynamic response to PD-1 blockade, suggesting common underlying immune principles with other ICB responsive tumor types. This immune-pharmacodynamic response alone, however, did not correlate with clinical responses. Thus, we developed a high-dimensional cytometry platform to examine multiple DDR pathways at single-cell resolution in subsets of human CD8 T cells relevant for response to ICB. This approach identified patterns of DDR in different subsets of CD8 T cells and defined a DDR signature associated with clinical outcome to PD-1 blockade. Specifically, we identified induction of phosphorylated ataxia telangiectasia mutated (ATM) (pATM) in CD8 T cells in the blood proliferating in response to anti-PD-1 in therapeutic responders but not in clinical non-responders. Collectively, these data identify T cell-intrinsic DDR and management of genome integrity during the proliferative response to PD-1 blockade as a potential biomarker of therapeutic benefit in uterine cancer patients. Moreover, the role of immune cell-intrinsic DDR may have implications in other contexts of immunotherapy and more broadly for human immune responses.

## RESULTS

### CD8 T cell pharmacodynamic response following PD-1 blockade in patients with hyper-mutated, chemotherapy-resistant uterine cancer

To test whether PD-1 blockade induced immune reinvigoration in patients with high TMB, we analyzed peripheral blood mononuclear cell (PBMC) from patients with advanced or recurrent MSI-H/MMRd or hypermutated uterine cancers treated with the anti-PD-1 antibody nivolumab (Clinicaltrials.gov NCT03241745, N = 35). All patients had previously received platinum-based chemotherapy and were eligible based on 1) MMR immunohistochemical testing or 2) evidence of MSI-H or TMB-H, defined as ≧20 mutations per megabase using the MSK integrated mutation profiling of actionable cancer targets (IMPACT) assay(29). Patients were treated with nivolumab, and PBMC examined every 2-4 weeks after initiating anti-PD-1 treatment. Detailed clinical outcomes for this cohort will be reported elsewhere (Friedman et. al, *In Preparation*). Clinical responders were defined based on RECIST criteria, with patients with complete response (CR) and partial response (PR) labeled as responders, and patients with stable disease (SD) or progressive disease (PD) labeled as non-responders. Tumors of responders exhibited a trend towards higher TMB in this cohort, consistent with previous studies (24–28) (**Supplementary Fig. 1A**). Other features, including patient age and days from the last chemotherapy to the initiation of nivolumab, did not differ between clinical response groups (**Supplementary Fig. 1B, C**). Because of the role of CD8 T cells in the response to PD-1 pathway blockade in other cancers (7–11), we focused on induction of the proliferation marker Ki67 in CD8 T cells in the peripheral blood. Indeed, in response to PD-1 blockade, a burst of CD8 T cell proliferation was observed 2-4 weeks after initiating treatment (**Fig. 1A, B**). This proliferative response returned to baseline by 8 weeks of treatment in most patients. We previously showed that in healthy donors, Ki67 expression by CD8 T cells varied only slightly over 3 weeks, changing 1.15-fold (range 0.88-1.69) (**Supplementary Fig. 2**)(9). Here, we used this baseline fluctuation in Ki67 in healthy untreated subjects to quantitatively define induction of a proliferative response following initiation of PD-1 blockade in this uterine cancer cohort. Most patients (21 out of 32 for whom PBMC cytometry was available) had a quantitative increase in Ki67 (> mean + 2 standard deviations (SD) of healthy donors) in CD8 T cells after initiation of anti-PD-1 treatment (**Fig. 1C, Supplementary Fig. 2**). Although the proliferative response to PD-1 blockade was observed in most patients, the magnitude of increase in %Ki67+ CD8 T cells after initiating anti-PD-1 treatment did not differ significantly between clinical responders versus non-responders based on RECIST criteria (**Fig. 1D**), suggesting other features of the immune response may influence the clinical outcome following immunotherapy.

**Fig. 1:**
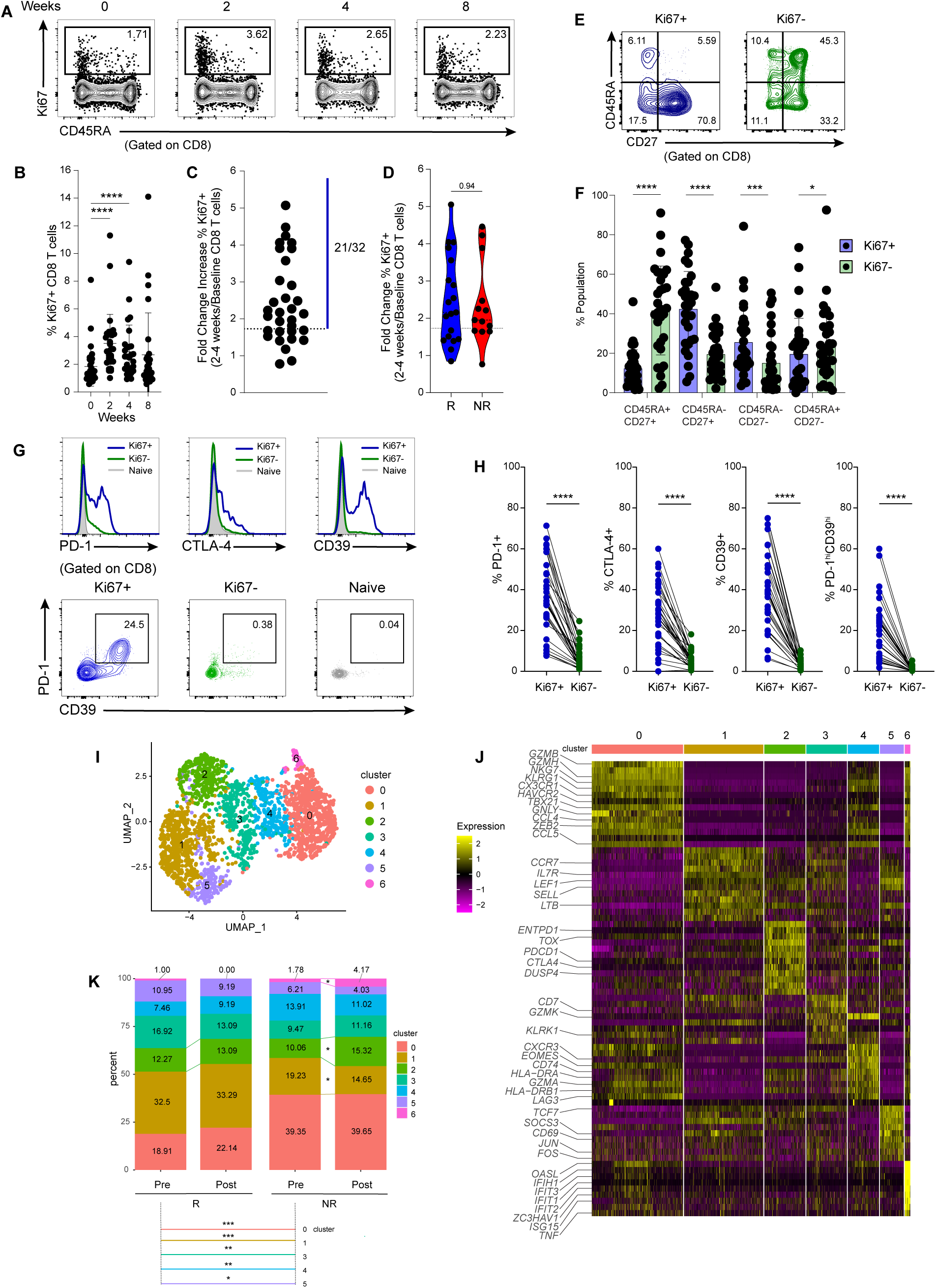
CD8 T cell response and kinetics of hyper-mutated chemotherapy resistant uterine cancer patients upon nivolumab treatment. Advanced or recurrent MSI-H/MMRd or hypermutated uterine cancer patients were treated with the anti-PD-1 antibody nivolumab. Blood was obtained before therapy, 2 weeks after therapy initiation and every 4 weeks thereafter on therapy. **(A, B)** Ki67 expression in CD8 T cells at indicated times (N = 32). **(C)** Fold change of Ki67 expression at peak of immunologic response versus pretreatment. Dotted line denotes fold change of 1.73, which is the mean plus 2 standard deviations (SD) of healthy donor CD8 T cell Ki67 variance over a similar time frame (see **Supplementary Fig. 2**). **(D)** Comparison of fold change of Ki67 expression at peak of immunologic response versus pretreatment between responders and non-responders (Responders (R: blue): N = 19, Non-Responders (NR: red): N = 13). Clinical response was determined by the best response by RECIST criteria. **(E)** Representative flow plots for Ki67+ (blue) and Ki67− (green) CD8 T cells at 2 weeks. **(F)**Distributions of indicated populations for Ki67+ (blue) and Ki67− (green) CD8 T cells at 2 weeks (N = 31). **(G)** Representative histogram flow plots for PD-1, CD39, CTLA-4 (top) and representative flow plots for PD-1^hi^CD39^hi^ Ki67+ (blue) and Ki67− (green) CD8 T cells (bottom) at 2 weeks post PD-1 blockade. **(H)** Expression of the indicated markers and populations of Ki67+ (blue) and Ki67− (green) CD8 T cells at 2 weeks (N = 31). **(I)** UMAP clustering of peripheral CD8 T cells from 8 individuals (5 responders and 3 non-responders) sampled pre-treatment and after 2 weeks of PD-1 blockade, across seven distinct Louvain clusters as represented by each color. **(J)** Heatmap showing distinct gene expression profiles for the identified clusters. Genes were selected from the 20 most significant differentially expressed genes across clusters plus genes of interest. **(K)** Cluster proportions across clinical response and time points at baseline (Pre) and at 2 weeks post initiation of nivolumab treatment (Post). * p <0.05, ** p <0.01, ***<0.001 by Fisher exact test. None of the comparisons reached FDR <0.05. Error bar denotes mean ± standard deviation (SD). For a violin plot (D), dashed line denotes median and dotted lines denote quantiles. * p <0.05, ** p <0.01, ***<0.001, **** < 0.0001 by unpaired Mann-Whitney U-test (D), Wilcoxon matched-pairs signed rank test (B, F, H).

To begin to examine the nature of these responding CD8 T cells, we used CD45RA and CD27 together with expression of inhibitory receptors to define T cell differentiation states. The responding Ki67+ CD8 T cells were largely CD45RA^lo^CD27^hi^ (**Fig. 1E, F**). Many of these responding CD8 T cells also had high expression of PD-1, CTLA-4, and/or CD39, and ∼20%-60% of this responding population was PD-1^hi^CD39^hi^ (**Fig. 1G, H**). These data are consistent with the phenotype of exhausted CD8 T cells (T_EX_) (8,9,30–33) and are in agreement with responses to PD-1 blockade in other cancers, where reinvigoration of T_EX_ is associated with the immune pharmacodynamic response to PD-1 immunotherapy (7–11).

To explore transcriptional dynamics of CD8 T cells responding to PD-1 blockade in uterine cancer patients, we performed single-cell RNA sequencing (scRNAseq) of non-naive CD3+ T cells (i.e. CD45RA+CD27+ “naive” T cells were excluded) sorted from PBMC from 5 responders and 3 non-responders, pre- and 2 weeks post initiation of anti-PD-1 treatment (see Methods). Although not possible to draw firm conclusions because of the limited number of patients with sufficient material for these paired analyses several observations from these data were of interest. There was little separation by time point, but a slight separation of responder and non-responder patients in the Uniform Manifold and Projection (UMAP) of these data (**Supplementary Fig. 3A**, **B**). Unsupervised clustering identified 7 distinct clusters of non-naive CD8 T cells (**Fig. 1I**). Clusters 0 and 4 were both more abundant in non-responders at baseline (p<0.01, 0.001 respectively), though these differences did not reach a false discovery rate (FDR) <0.05, in part because of a low number of patients for whom samples were sufficient for scRNA-seq analysis (**Figure 1K, Supplementary Fig. 3B**). Cluster 0 had high expression of effector-associated genes such as *GZMB, GLNY, KLRG1, CX3CR1, TBX21, HAVCR2* and *ZEB2*, and was more abundant in clinical non-responders at both time points (**Fig. 1I-K, Supplementary Fig. 3B-D**). Cluster 4 had high expression of MHC Class II genes including *HLA*-*DRA* and *HLA-DRB1* as well as CD74, and also expressed *EOMES* and *CXCR3* (**Fig. 1J, K, Supplementary Fig. 3C, D**). Cluster 6 was a minor cluster defined by high expression of interferon-stimulated genes (ISGs) including *OASL*, *ISG15* and several *IFIT* genes, as well as high expression of *TNF*. This ISG-enriched cluster was present both in clinical responder and non-responders at baseline, but more abundant and increased >2-fold post PD-1 blockade in a non-responder (**Fig. 1J, K, Supplementary Fig. 3B-D**). In contrast, clusters 1, 3 and 5 were more abundant in responders at baseline based on p value (p <0.001, 0.01, 0.05 respectively) though did not reach FDR<0.05 (**Fig. 1K, Supplementary Fig. 3B**). Cluster 1 had high expression of genes such as *CCR7, IL7R*, and *SELL* (encoding CD62L) associated with central memory (**Fig. 1J, K, Supplementary Fig. 3 C, D**). This cluster was more abundant in clinical responders at baseline, and decreased post treatment in some clinical non-responders (**Fig. 1I-K, Supplementary Fig. 3 B-D**). Cluster 3 had high expression of *CD7* and *KLRK1* and Cluster 5 had high expression of *TCF7* and *SOCS3,* as well as genes related to TCR signaling including *CD69, JUN* and *FOS* (**Figure 1J, K**) Cluster 5 also shared expression of some genes such as *IL7R* and *SELL* with cluster 1. Cluster 2 had characteristics consistent with T_EX_ cells, including high expression of *TOX*, *PDCD1*, *CTLA4* and *ENTPD1* (encoding CD39) (**Fig. 1J, Supplementary Fig. 3C, D**). This cluster showed a trend to increase in clinical responders, but also increased in one non-responder, with some variation across patients (**Fig. 1J, K, Supplementary Fig. 3B, E**), in agreement with the flow cytometry data above and previous data for other cancers(7–11). These scRNA-seq data also highlight that clinical non-responders had roughly double the size of an effector-like CD8 T cell cluster (cluster 0). In contrast, clinical responders had more central memory and fewer effector-like CD8 T cells. Thus, in addition to highlighting some patient-to-patient variation, this scRNA-seq analysis suggested a different balance of effector/effector memory-like and central memory-like CD8 T cells in clinical responder and non-responder. The data also resolved T_EX_-like cells in both patient groups and was consistent with the flow cytometry. This scRNAseq data collectively indicate that numerical changes in CD8 T cell subsets or transcriptionally defined CD8 T cell clusters did not easily distinguish clinical responder and non-responder patients.

Overall, these data are consistent with an immunological response to anti-PD-1 treatment in MSI-H/MMRd or hypermutated uterine cancer patients that involves reinvigoration of T_EX_. However, these pharmacodynamic responses alone did not explain differences in clinical outcomes.

### Development and validation of a DDR-Immune profiling platform

A major gap in our understanding of ICB for cancer is why reinvigoration of T cells alone does not fully explain clinical responses. These observations suggest additional, unknown, immunological determinants of beneficial response. Key to the immunological response to PD-1 blockade is reinvigorating the proliferative response of T_EX_ (1, 2) and/or induction of new T cell responses, likely effector-like T cell subsets. Recent work has highlighted the importance of DNA damage and genome integrity in CD8 T cells during antigen-driven proliferation (20), but the role of cell-intrinsic DDR in CD8 T cells in patients on checkpoint blockade has not been examined. Because most patients mounted a proliferative response to PD-1 blockade (perhaps because of the generally high TMB), we hypothesized that T cell-intrinsic DDR might help explain the disconnect between immunological and clinical responses.

To dissect the role of T cell-intrinsic DDR, we developed a flow cytometry-based DDR-immune profiling platform. This approach allowed us to simultaneously interrogate T cell differentiation state, together with changes in major DDR pathways (**Fig. 2A**). We developed approaches to examine markers of double strand break (DSB) repair, homologous recombination (HR), non-homologous end joining (NHEJ), base excision repair (BER), mismatch repair (MMR), and nucleotide excision repair (NER) (34, 35) at the single cell level by flow cytometry where we could also interrogate other features of T cell differentiation states (**Fig 2A**).

**Fig. 2:**
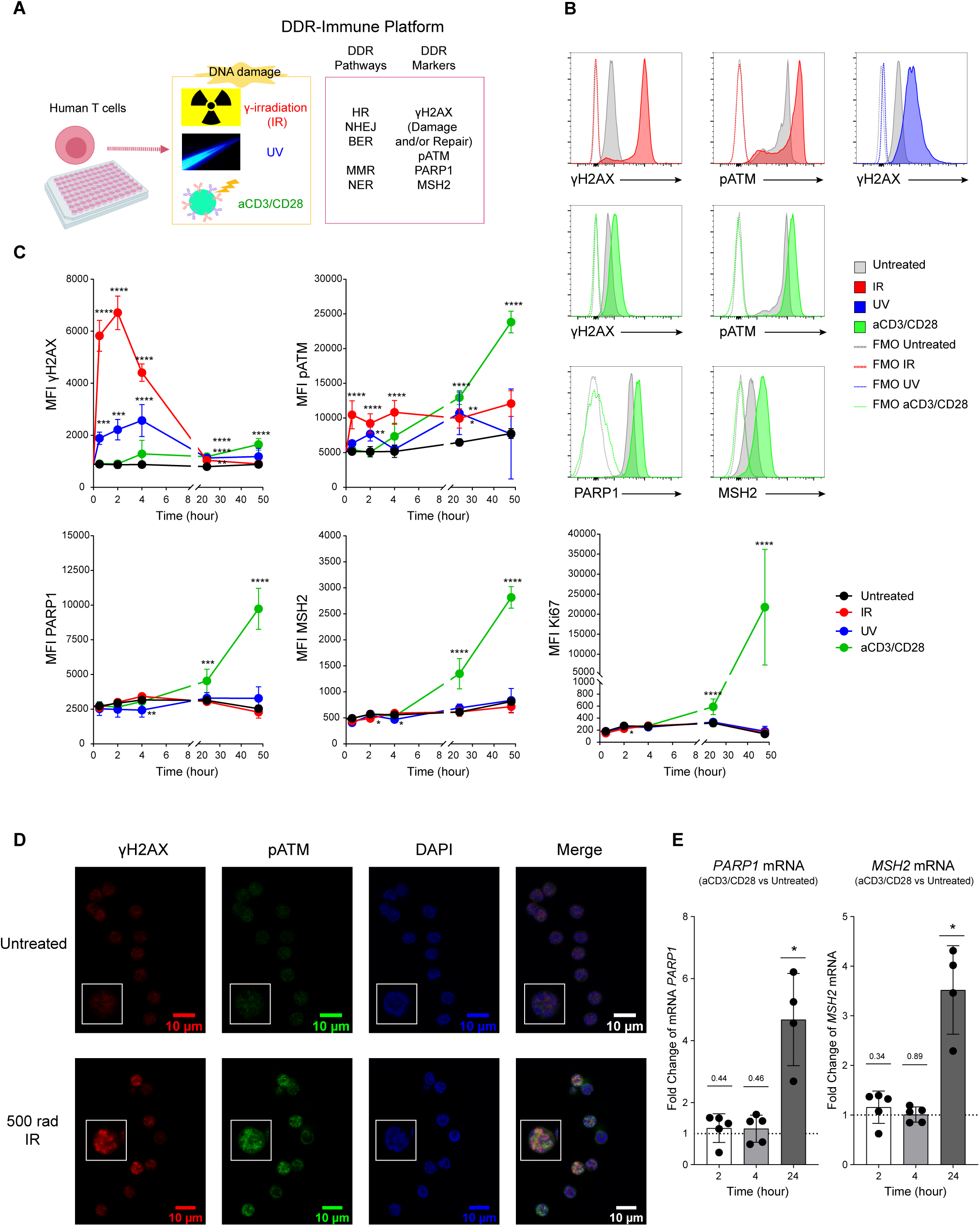
CD8 T cell responses to DNA damage. **(A)** Overview of DDR-immune platform. Human T cells isolated from PBMC were treated with γ-irradiation (IR), UV, or stimulated with aCD3/CD28. DDR markers and relevant pathways listed here (gating strategies in **Supplementary Fig. 4**). Illustration created with BioRender.com. **(B)** DDR of human CD8 T cells. CD8 T cells were treated with IR, UV, aCD3/CD28 and incubated for 2 hours for IR, 4 hours for UV and 24 hours for aCD3/CD28. DDR makers were assessed by flow cytometry. Representative histogram flow plots. **(C)** Time kinetics of mean fluorescence intensity (MFI) of DDR markers (γH2AX, pATM, PARP1, MSH2) and proliferation marker (Ki67), upon different types of DNA damage. **(D)** Representative Immunofluorescence (IF) image with magnified view (inset) of irradiated/untreated human CD8 T cells stained for γH2AX and pATM. Scale bars: 10 μm. **(E)** Representative mRNA expression of *PARP1* and *MSH2* post aCD3/CD28 stimulation. CD8 T cells were stimulated with aCD3/CD28 and 2, 4 and 24 hours later, cells were examined for gene expression using qPCR. (C) N = 5, Error bars: standard deviation (SD). * p <0.05, ** p <0.01, *** p<0.001, **** p<0.0001 by one-way ANOVA with Dunnett’s test, compared with untreated T cells. (E) N = 3 (samples), 3 technical replicates per sample were averaged. Error bar denotes mean ± SD. * p<0.05 by one-sample t test. Abbreviations: Homologous recombination (HR), non-homologous end joining (NHEJ), base excision repair (BER), mismatch repair (MMR), and nucleotide excision repair (NER).

We first used this platform to examine T cells from healthy subjects exposed to different types of DNA damage. We treated T cells from healthy donors with γ-irradiation (IR), ultraviolet light (UV) or anti-CD3/CD28 beads (“aCD3/CD28”) (see Methods). IR mainly induces DSB, whereas UV preferentially induces single-strand breaks (SSB) and pyrimidine dimers (34). Stimulation with anti-CD3/CD28 (aCD3/CD28) beads leads to robust T cell proliferation, modeling “proliferative and replication stress”, likely at replication forks and telomeres (34, 36). We first examined the kinetics of induction of different DDR pathways in CD8 T cells (**Fig. 2B, C**). DNA damage induces autophosphorylation of Phosphoinositide-3-Kinase (PI3K)-like Kinase (PIKK) ataxia telangiectasia mutated (ATM) on serine 1981 causing dimer dissociation and activation of the ATM kinase (37). Upon recognizing double strand DNA breaks, ATM phosphorylates the histone variant H2AX on serine 139 (called γH2AX) and γH2AX then serves as a scaffold for proteins to initiate repair of DNA damage (38). Indeed, upon induction of DNA breaks by IR, γH2AX was rapidly induced, peaking at ∼2 hours and returning to the baseline by 24 hours. IR also induced phosphorylation of ATM (pATM) as early as 30 minutes, consistent with the γH2AX observation (**Fig. 2B, C**). This pATM signal was sustained for at least 48 hours. UV irradiation also induced γH2AX, but the peak of this UV-induced signal was later than following IR and the overall magnitude was lower (**Fig. 2B, C**). pATM was not only induced upon IR together with γH2AX, but also increased following aCD3/CD28 stimulation with less robust induction of γH2AX at early time points, but gradual accumulation of this DDR response marker over time (**Fig. 2B, C**), consistent with a role for pATM after T cell receptor (TCR) stimulation (39). PARP1 and MSH2 are involved in base excision repair and mismatch repair (34) and are of interest, at least partly, because of the widespread use of PARP inhibitors in immune oncology treatment combinations (35) and the germline mutations in MSH2 in some Lynch patients who have hypermutated cancers(40). PARP1 and MSH2 expression were both increased at 24-48 hours after TCR stimulation, consistent with their roles in replication-coupled DNA damage repair (34, 36) (**Fig. 2B, C**). The kinetic differences in γH2AX and pATM signals versus PARP1 and MSH2 also may reflect the rapid post-translational modification for the former (γH2AX and pATM) versus the likely need for new protein expression for the latter (PARP1 and MSH2).

To complement the flow cytometric approach and further validate the specificity of this platform, we performed immunofluorescence and quantitative Real-Time Reverse Transcription PCR (qRT-PCR). Indeed, increased nuclear staining of γH2AX and pATM upon IR was observed by immunofluorescence (IF) (**Fig. 2D**), confirming the flow cytometric approach above. Increased expression of *MSH2* and *PARP1* upon aCD3/CD28 stimulation was also validated by qRT-PCR (**Fig. 2E**). Thus, this DDR-immune platform allowed robust detection of the multiple DDR pathways in human T cells upon introduction of DNA damage and proliferative stress.

### Distinct DDR patterns and sensitivity to DNA damage in different CD8 T cell subsets

The response to external signals can vary for CD8 T cells in different states of differentiation. For example, responses to cytokines (41–45) or chemokines (46, 47) can be distinct in naive, effector, effector memory, or central memory T cells. However, the impact of DNA damage and the response of DDR pathways in different human CD8 T cell subsets have not been defined. Moreover, it is becoming increasingly clear that individual subtypes of human T cells (e.g. T_EX_, Stem Cell Memory) could be preferentially important for response to immunotherapy. We therefore next examined DDR in 7 distinct human CD8 T cell subsets: Naive CD8 T cells (T_Naive_: CD45RA+CD27+CCR7+CD95-), Stem Cell Memory (T_SCM_: CD45RA+CD27+CCR7+CD95+), Central Memory (T_CM_: CD45RA-CD27+CCR7+), two different Effector Memory populations distinguished by CD27 expression (T_EM1_: CD45RA-CD27-CCR7- and T_EM2_: CD45RA-CD27+CCR7-), Terminally differentiated Effector Memory RA (T_EMRA_: CD45RA+CD27-CCR7-), and Exhausted (T_EX_ : non-naive PD-1^hi^CD39^hi^) CD8 T cells (31,48–53) (**Supplementary Fig. 4**). We applied the DDR-immune profiling approach described above to these CD8 T cell subsets from healthy donors. CD8 T cells were treated with IR, UV, aCD3/CD28 and incubated for 2 hours for IR, 4 hours for UV and 24 hours for aCD3/CD28 based on the kinetics defined above and induction of these DDR markers was examined by flow cytometry for each CD8 T cell subset (**Fig. 3A**, **Supplementary Fig. 5**). This DDR-Immune platform revealed CD8 T cell subset-specific patterns of DDR, as well as specific DDR pathways induced by different types of DNA damage. Notably, T_SCM_ and T_EMRA_ had higher γH2AX than other T cell subsets especially following IR-induced damage (**Fig. 3A-D, Supplementary Fig. 5A**). Elevated γH2AX could indicate an increase in DNA damage and loss of genome integrity or an increased response to damage and induction of repair. Whereas increased DNA damage would lead to increased cell death, repair of this genomic damage may not. To distinguish these possibilities, we examined activated caspase 3 in combination with viability staining (Live/Dead (L/D)) in each subset before and after introducing DNA damage (**Supplementary Fig. 6 A-C**). Before IR T_EM1_ and T_EX_ had significantly higher active caspase 3 and L/D staining than other populations. After introducing DNA damage by IR, however, caspase 3 increased robustly in T_EM1_ and T_EMRA_ populations, modestly increased in T_CM,_ T_EM2_ and T_EX_, and remained low in T_Naive_ and T_SCM_ (**Supplementary Fig.6 A-C**). These data suggested that introduction of DNA damage was associated with poor survival and increased cell death in more differentiated CD8 T cell subsets (T_EM1_ and T_EMRA_). However, the increase in γH2AX in T_SCM_ but low induction of caspase 3 and L/D staining suggested that the γH2AX increase in this population was likely connected to efficient DNA repair, consistent with recent studies in mice(20). Indeed, the ratio of γH2AX to caspase 3 was substantially higher in T_SCM_ compared to other populations (**Supplementary Fig. 6D**). In contrast, T_EX_, TEMRA and especially T_EM1_ had the lowest ratio of γH2AX to caspase 3 (**Supplementary Fig. 6D**). A similar trend was observed for pATM upon IR. Though pATM expression was highest in T_EMRA_, the ratio of pATM to caspase 3 was substantially higher in T_SCM_ compared to other populations, whereas T_EX_, T_EMRA_ and especially T_EM1_ had the lowest ratio (**Fig.3 A, C**, **Supplementary Fig. 5B, Supplementary Fig. 6E**). Following UV-mediated DNA damage, induction of γH2AX and pATM was modest and occurred in all CD8 T cell subsets, with higher induction in T_SCM_ and T_CM_ (**Fig. 3A, C, D, Supplementary Fig. 5C, D**). In contrast, PARP1 and MSH2 were not induced by UV or IR (**Fig. 3C, D**). A different pattern was observed after aCD3/CD28 stimulation. Cell division, as indicated by Ki67, was increased in all CD8 T cell subsets with the highest Ki67 induction in T_SCM_, T_CM_, T_EM2_ and T_EX_ cells followed by T_EM1_, T_EMRA_ and T_Naive_ (**Figs. 3C, D**). Although examined at an early timepoint, it is possible that aCD3/CD28 stimulation causes some T_Naive_ to become activated to fall outside of the strict naive gates used in this assay because of loss of naive phenotypic markers. Nevertheless, during proliferative stress, although γH2AX was low in all subsets (perhaps reflecting the time point examined), pATM was induced and most highly upregulated in T_EX_, followed by T_CM_ and then T_SCM_ and T_EM2_, with T_EM1_ and T_EMRA_ displaying lower pATM, only modestly above T_Naive_ (**Fig. 3C, D**). A similar hierarchical pattern was observed for MSH2 and PARP1 following proliferative stress with upregulation in T_EX_, T_SCM_, T_CM_ (and T_EM2_ for PARP1), followed by T_EMRA_, (**Fig. 3C, D, Supplementary Fig. 5E, F**). The association of MSH2 and PARP1 with proliferation was consistent with the association of MMR and base excision repair with DNA repair upon replication (34, 36). Collectively, these data illustrate distinct DDR patterns among different T cell subsets, patterns that could contribute to different susceptibility to genomic damage and cell death.

**Fig. 3:**
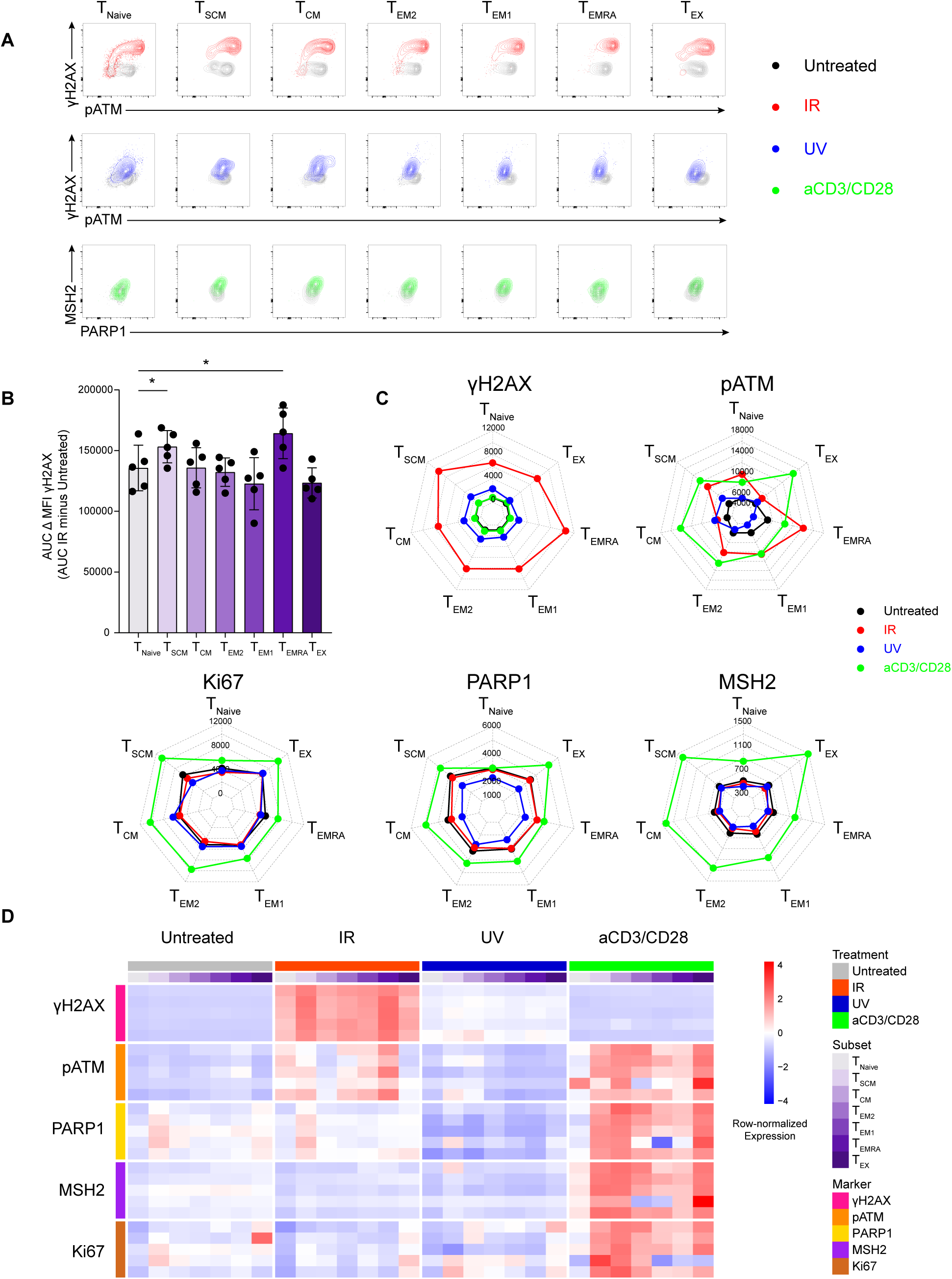
CD8 T cell subsets have distinct DDR patterns. Human T cells were treated as described in Fig. 2 and Methods, and cell populations were gated as described in **Supplementary ig. 4. (A)** Representative flow cytometry plots for DDR markers. **(B)** Difference (Δ) between the untreated and irradiation-induced γH2AX MFI Area under curve (AUC) in CD8 T cells through 24 hours post treatment. **(C)** Radar plots of DDR markers across different CD8 T cell subsets. The values are median of MFI for each marker. **(D)** Heatmap of DDR markers across T cell subsets and DNA damage treatments. (B) N = 5, Error bar denotes mean ± SD. * p <0.05 by one-way ANOVA with Dunnett’s test, compared with naive T cells.

### Clinical Response to ICB is associated with distinct CD8 T cell-intrinsic DDR signatures

The data above indicate that different CD8 T cell subsets engage distinct patterns of DDR depending on the type of DNA damage stress induced. These observations provoke the question of whether DDR patterns are associated with proliferative responses induced by anti-PD-1 treatment in human cancer patients. The MSI-H or hypermutated uterine cancer cohort examined here is of potential interest given the potential high antigen burden in these patients that could drive T cell proliferation following ICB. In addition, these patients received previous treatment with DNA damaging chemotherapy and/or pelvic radiation that may affect immune cells, in addition to cancer cells.

In this uterine cancer cohort, the fold change in Ki67 in responding CD8 T cells after the first treatment dose of anti-PD-1 did not distinguish clinical responders and non-responders (**Fig. 1D**). Indeed, there was also no difference in the timing, kinetics, or durability of this proliferative response following PD-1 blockade between clinical responders and non-responders (**Fig. 4A**). To examine whether CD8 T cell-intrinsic DDR responses in the responding, proliferating CD8 T cells might distinguish patients who clinically respond to PD-1 blockade versus those who do not, we applied the DDR-Immune profiling platform to peripheral blood CD8 T cells in these MSI-H endometrial cancer patients receiving PD-1 blockade. Overall, Ki67+ CD8 T cells, most of which were T_EX_ (**Fig. 1**), expressed higher pATM, compared with Ki67-CD8 T cells at baseline and at 2 weeks after initiation of anti-PD-1 treatment (**Fig. 4B, Supplementary Fig. 7C, D**). Ki67+ CD8 T cells also had higher expression of other DDR markers, γH2AX, PARP1 and MSH2 at 2 weeks (**Supplementary Fig. 7A, B**) and at baseline (**Supplementary Fig. 7C, D**). We extended these analyses to examine how these DDR pathways related to markers of CD8 T cell activation and exhaustion within the responding Ki67+ population at baseline, at the peak of the proliferative response at 2-4 weeks, and at 8 weeks after initiation of treatment when proliferative responses had returned to baseline. At baseline, high expression of pATM was positively correlated with CD28, and negatively associated with expression of TBET and EOMES (**Fig. 4C**). At the peak of the proliferative response 2 weeks after initiating anti-PD-1 treatment, expression of pATM was strongly correlated with PD-1 (FDR <0.05) (**Fig. 4D**). At 4 weeks, pATM was still positively associated with markers of T_EX_, including PD1, CD39, CD69 and CTLA-4, as well as CD28 (**Fig. 4E**), and the trend continued at 8-week time points (**Fig. 4F**). The most statistically robust of these relationships was the correlation of pATM with PD-1 expression at week 2. Other associations also support the notion of a DDR response in responding CD8 T cell populations, but did not achieve FDR<0.05, perhaps because of heterogeneity in the CD8 T cell-intrinsic DDR response across patients. Nevertheless, these data support a potential role for CD8 T cell-intrinsic pATM pathway engagement in T_EX_ phenotype CD8 T cells upon reinvigoration by PD-1 blockade.

**Fig. 4:**
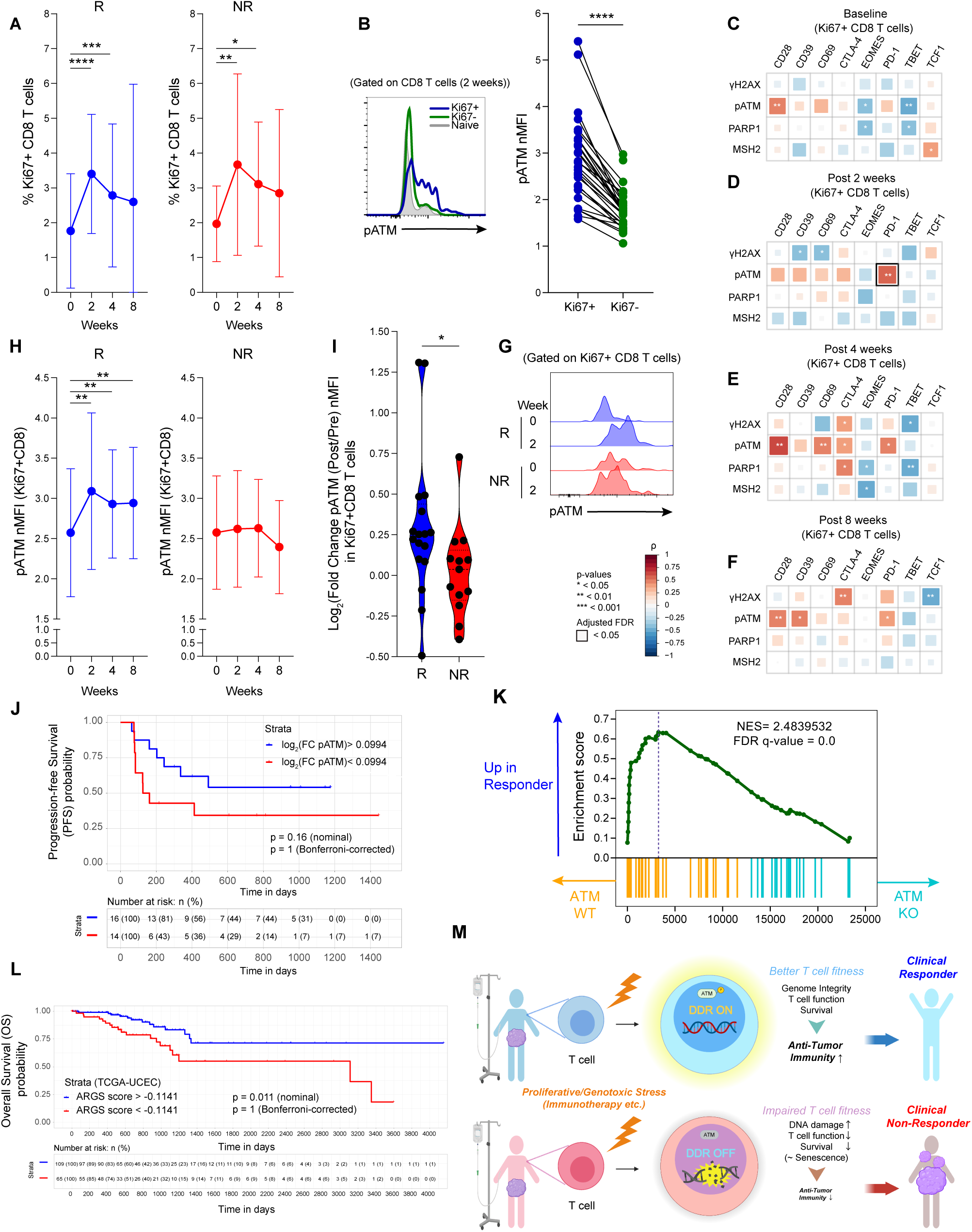
CD8 T cell-intrinsic DDR is an independent variable associated with clinical responses to PD-1 blockade in MSI-H endometrial cancer. **(A)** Ki67 expression in CD8 T cell subsets upon nivolumab treatment at indicated times (Responders (R: blue): N = 19, Non-Responders (NR: red): N = 13). * p <0.05, ** p <0.01, *** p<0.001, **** p<0.0001 by two-sided Wilcoxon matched-pairs test. Error bar denotes mean ± SD. **(B)** Left: Representative flow plot for pATM expression; gated on Ki67+ CD8 (blue), Ki67-CD8 (green) and Naive CD8 (gray). Right: Normalized MFI (nMFI) of pATM in Ki67+ (blue) and Ki67− (green) CD8 T cells at 2 weeks post nivolumab treatment (N = 30). **** < 0.0001 by Wilcoxon matched-pairs signed rank test. MFI values were normalized to those of naive CD8 T cells of healthy donor control. **(C-F)** Correlation matrix of DDR markers and immune features in Ki67+ CD8 T cells from baseline **(C)**, 2 weeks post nivolumab treatment **(D)**, 4 weeks post nivolumab treatment **(E)**, and 8 weeks post nivolumab treatment time points **(F)**. Immunological marker expression is shown as % positivity of Ki67+ CD8 T cells. DDR features are shown as normalized MFIs (nMFI), in which MFI values were normalized to those of naive CD8 T cells of healthy donor control. Spearman’s Rank Correlation coefficient (ρ) is indicated by square size and heat scale; significance indicated by: * p<0.05, ** p<0.01, and *** p<0.001; bold black box indicates FDR<0.05. **(G)** Representative flow plots for pATM expression of pre-versus 2 weeks post-initiation of nivolumab treatment, gated on Ki67+ CD8 in responder (R) and non-responder (NR). **(H)** Patterns of pATM expression kinetics in CD8 T cell in Responder versus Non-Responder (Responders (R: blue): N = 17, Non-Responders (NR: red): N = 13); ** p <0.01 by two-sided Wilcoxon matched-pairs test. Error bar denotes mean ± SD. **(I)** Log2 fold change increase (FC) of pATM from baseline to 2 weeks in Ki67+ CD8 T cells. (Responders (R: blue): N = 17, Non-Responders (NR: red): N = 13). * p <0.05 by unpaired Mann-Whitney U-test. Dashed lines denote median, and dotted lines denote quantiles. **(J)** Kaplan-Meier plot of the Progression Free Survival (PFS) between high (>0.0994) versus low (< 0.0994) log fold change (FC) increase of pATM from baseline to 2 weeks in Ki67+ CD8 T cells (Log_2_(FC pATM)). Blue line indicates Log_2_(FC pATM) > 0.0994, red line indicates Log_2_(FC pATM) < 0.0994. Cut off was determined by *cutpointr* (see Material and Methods). Nominal p-value was determined by log-rank test (p = 0.16, Bonferroni adjusted p= 1). **(K)** Gene Set Enrichment Analysis (GSEA) showed enrichment of responder gene signature in proliferating CD8 T cells derived from scRNAseq (see methods for detail) was analyzed in transcriptomic data from ATM^WT^ versus ATM^KO^ human CD8 T cells. FDR and normalized enrichment score (NES) are indicated. Dotted line indicates the leading edge. **(L)** Kaplan-Meier plot of the Overall Survival (OS) between high (> −0.1141) versus low (< −0.1141) ATM-Responder gene signature (ARGS) score by Gene Set Variation Analysis (GSVA) of RNAseq2 expression data from The Cancer Genome Atlas Uterine Corpus Endometrial Carcinoma (TCGA-UCEC) data collection (N = 174). Blue line indicates ARGS score > −0.1141, red line indicates ARGS score < −0.1141. Cut off was determined by *cutpointr* (see Material and Methods). Nominal p-value was determined by log-rank test (p = 0.011, Bonferroni adjusted p= 1). **(M)** Graphical summary of the model. Illustration created with BioRender.com.

We next examined the relationship between DDR and immune markers in proliferating T cells upon PD-1 blockade with clinical parameters. Age, TMB and days since prior chemotherapy before starting nivolumab, were not significantly correlated with the DDR response in proliferating CD8 T cells. However, a positive correlation was observed between TMB and CD69 and also between days since prior chemotherapy and TCF1 in Ki67+ CD8 T cells at the peak of the proliferative response to PD-1 blockade, though these correlations did not reach FDR <0.05 (**Supplementary Fig. 8**).

In this cohort, 5 patients had germline mutation of MMR genes (*MSH2* mutation (N = 3), *PMS2* mutation (N = 1), and germline *MLH1* epimutation (N = 1)). To begin to investigate whether germline mutations in DNA repair pathways influenced the induction of DDR in CD8 T cells responding to PD-1 blockade, we compared patients with and without germline MMRd (gMMRd). Overall, patients with gMMRd tended to have higher TMB and were younger (**Supplementary Fig. 9A, B**), but the duration since last chemotherapy in those with and without gMMRd was similar (**Supplementary Fig. 9C**). We hypothesized that gMMRd patients might exhibit a different pattern of proliferative response to anti-PD-1 and/or T cell-intrinsic DDR given the DNA repair deficiencies due to mutations in these genes in T cells as well as cancer cells. However, the induction and kinetics of Ki67 in responding CD8 T cells were compatible between gMMRd and non-gMMRd patients (**Supplementary Fig. 9D-G**). Furthermore, frequency of T_EX_ (PD-1^hi^CD39^hi^ in non-naive CD8 T cells) pre and post PD-1 blockade was comparable, albeit the number of gMMRd patients is small (**Supplementary Fig. 9H, I**). A limited sample size of only 5 gMMRd (spanning 3 different genes mutated) might not provide sufficient resolution of an effect of germline DDR mutations in this setting and studies in larger cohorts of gMMRd patients are likely warranted to further delineate the impact of gMMRd on T cells and immune responsiveness.

To investigate how changes in pATM in Ki67+ CD8 T cells responding to PD-1 blockade were related to clinical outcome, we examined pATM in Ki67+ CD8 T cells in clinical responders versus non-responders after initiating PD-1 blockade therapy. In clinical responders there was a robust induction of pATM in Ki67+ CD8 T cells at 2 weeks after initiation of PD-1 blockade (**Fig. 4G, H**). This pATM induction in clinical responders contrasted with a near absence of pATM induction in Ki67+ CD8 T cells in clinical non-responders at 2 weeks after starting anti-PD-1 (**Fig. 4G, H**). As an additional approach, we examined the log 2-fold change (log FC) increase of pATM (log_2_ (FC pATM)) from baseline to 2 weeks in Ki67+ CD8 T cells. Indeed, clinical responders had higher log FC increase in pATM induction upon PD-1 blockade, compared to non-responders who, on average had no pATM induction in Ki67+ CD8 T cells upon therapy initiation (**Fig. 4I**). We also examined other DDR features. No clear trends were observed in γH2AX expression at the time points analyzed (**Supplementary Fig. 10 A, D**). PARP1 expression remained unchanged or decreased in some clinical non-responders at 2 weeks but remained unchanged or increase in a subset of clinical responders, though the magnitude of change in PARP1 was less robust than pATM (**Supplementary Fig. 10B, E**). Changes in MSH2 were relatively modest in both clinical responders and non-responders (**Supplementary Fig. 10C, F**). Thus, induction of DDR, especially changes in pATM, in CD8 T cells responding to checkpoint blockade was a feature of clinical responding patients in this uterine cancer cohort.

Next, we investigated how pATM induction related to the co-primary endpoint of progression-free survival (PFS). We separated patients into those with high (log_2_ (FC pATM) > 0.0094) versus low (log_2_ (FC pATM) < 0.0094) induction of pATM in Ki67+ CD8 T cells responding to PD-1 blockade (see Methods). Patients with higher induction of pATM in responding Ki67+ CD8 T cells at the peak of the immunological response to PD-1 blockade trended to having longer PFS (**Fig. 4J**). This cohort was not initially designed for this analysis since this DDR signature was only discovered by deeply profiling these patients and this result did not reach statistical significance (nominal p-value = 0.16, Bonferroni-adjusted p-value =1) (**Fig. 4J**). We estimated that nominal significance would require N = 65 patients or more to be appropriately powered (see Methods) and this number of patients was not available on this current trial. We therefore turned to two complementary approaches to further address this DDR biology in immunotherapy outcomes.

First, we examined whether there was a relationship between transcriptional responses to ATM activation in T cells responding to therapy in clinical responder versus non-responders. To test this idea, we first generated a transcriptional signature of ATM-dependent biology using primary human T cells that rendered ATM deficient by CRISPR-mediated knockout (KO) (see Methods and **Supplementary Fig. 11**). These ATM^KO^ human T cells were viable and capable of proliferating *in vitro* with population doublings and cell volume changes during T cell culture comparable to control WT T cells treated with a control non-targeting sgRNA (**Supplementary Fig. 11A**). These ATM^KO^ T cells had an indel frequency of ∼80% indicating a high efficiency of deletion of *ATM* (**Supplementary Fig. 11B**). Western blot confirmed diminished expression of total and phosphorylated ATM in ATM^KO^ CD8 T cells compared with control targeted CD8 T cells (**Supplementary Fig. 11C-F**). Induction of DNA damage by irradiation also failed to upregulate pATM in ATM^KO^ T cells (**Supplementary Fig. 11G, H**). We used these ATM^KO^ CD8 T cells to generate an ATM-dependent transcriptional signature of the response to proliferation. RNAseq identified a set of ATM-dependent genes by comparing ATM^WT^ and ATM^KO^ CD8 T cells (**Supplementary Table 2**). This proliferating CD8 T cell ATM DDR signature was used to test whether this signature was enriched in Ki67+CD8 T cells from responder versus non-responder patients. Thus, we generated transcriptional signatures of CD8 T cells responding to anti-PD-1 in clinical responder versus non-responders (FDR <0.05) by examining differentially expressed genes from the scRNA-seq analysis specifically in proliferating CD8 T cells (in the S and/or G2M phase of the cell cycle) after initiating PD-1 blockade (**Supplementary Table 3**, **Supplementary Fig. 12**). We then tested whether the ATM-dependent transcriptional signature was differentially enriched in proliferating CD8 T cells from responder versus non-responder patients. Indeed, CD8 T cells proliferating in response to anti-PD-1 therapy from clinical responders strongly enriched in the transcriptional signature of intact ATM activity (i.e. transcriptional signature up in ATM^WT^ compared to ATM^KO^) compared to clinical non-responders (**Figure 4K**). These data support notion that ATM regulated transcriptional circuits in CD8 T cells are engaged upon anti-PD-1 treatment and that the ability to induce this ATM-dependent transcriptional response to DNA damage was associated with better clinical response to PD-1 blockade.

Finally, we investigated the association between the ATM regulated gene networks in T cells responding to PD-1 blockade and survival outcome using the The Cancer Genome Atlas (TCGA) datasets. We first examined data for Uterine Corpus Endometrial Carcinoma (UCEC) patients. We used leading-edge genes from the GSEA analysis above as an “ATM-Responder Gene Signature (ARGS)” in T cells (**Supplementary Table 4**), and calculated an ARGS score by Gene Set Variation Analysis (GSVA)(54). We then separated patients into those with high versus low ARGS scores (see Methods). In this UCEC cohort (N=174), patients with higher ARGS score (high: ARGS score > - 0.1141, low: ARGS score < −0.1141) exhibited longer overall survival (OS), although this association did not meet significance after Bonferroni-adjustment (nominal p-value = 0.011, Bonferroni-adjusted p-value =1) (**Fig. 4L**). Subclass analysis of MSI-H UCEC (N =46) also revealed a similar trend (**Supplementary Fig. 13A**). Of course, the major contributor to these signatures in TCGA data will be tumor cells and stromal cells rather than immune cells. To begin to examine if the ARGS was associated with CD8 T cells in the TCGA dataset. We compared the ARGS score between patients with high *CD8A* expression versus low *CD8A* expression. Indeed, there was significantly higher expression of ARGS in the high *CD8A* group (**Supplementary Fig. 13B**). We next examined whether these observations could be extended to other cancers. Indeed, in colon adenocarcinoma (COAD), patients with higher ARGS score (high: ARGS score > −0.1066, low: ARGS score < −0.1066) exhibited longer OS by nominal p-value (nominal p-value = 0.028, Bonferroni-adjusted p-value =1) (**Supplementary Fig. 13C**). There were also similar trends in the subset of MSI-H COAD patients and skin cutaneous melanoma (SKCM), though smaller sample size limited statistical power for these datasets for OS analyses (**Supplementary Fig. 13D, F**). Nevertheless, in COAD and SKCM datasets, the ARGS score was significantly higher in patients with high *CD8A* expression (**Supplementary Fig. 13E, G**). Thus, although some of the enrichment in the TCGA datasets could reflect differences in DDR in cancer cells or other non-immune cells, these data are consistent the notion that a transcriptional circuit induced by ATM in CD8 T cells may be associated with favorable clinical outcomes in different types of cancers.

Collectively, these data provoke a model where CD8 T cell-intrinsic DDR pathways, and specifically induction of pATM in Ki67+ T_EX_-like CD8 T cells reinvigorated by PD-1 blockade have a key role in beneficial clinical outcome to immunotherapy. These data also identify CD8 T cell-intrinsic DDR as a potential immunological signature of clinical response to immune checkpoint blockade (**Fig. 4 M**).

## DISCUSSION

Despite the benefit of PD-1 blockade in many cancer patients, a substantial proportion do not derive long-term benefit. In this study, we examined the relationship between immunological response to PD-1 blockade and clinical outcomes in a cohort of advanced or recurrent MSI-H or hypermutated uterine cancer patients. Pembrolizumab is FDA-approved for the treatment of MSI-H and hypermutated advanced solid tumors (21,22,55). The MSI-H or hypermutated uterine cancer cohort examined here was of interest because this patient group has generally high TMB, but the immunological associations with outcome of PD-1 blockade remain poorly understood. Even in these patients where there is often high tumor mutational burden optimal clinical benefit is not always achieved. We identified an immune pharmacodynamic response to PD-1 blockade in the blood that included induction of proliferation of T_EX_ phenotype CD8 T cells, adding uterine cancer to a number of cancer types where T_EX_ are a major cell type responding to PD-1 pathway blockade(7–11). This immune response to PD-1 blockade, however, did not correlate with clinical benefit, suggesting other immune features associated with disease control. Indeed, we identified a previously unappreciated CD8 T cell-intrinsic response to DNA damage in CD8 T cells responding to PD-1 blockade that distinguished clinical responders from non-responders. Thus, these studies revealed several key findings. First, in this cohort, the majority of patients responded immunologically to anti-PD-1 treatment indicating an “on target” effect for reinvigorating pre-existing T_EX_ and/or inducing new T cell responses. Second, we defined signatures of response to different types of DNA damage in human CD8 T cells.

These signatures differed depending on CD8 T cell subset indicating potential relevance of the responding T cell type for immunotherapy of cancer. Third, we identified a DDR signature in CD8 T cells responding to anti-PD-1 therapy that distinguished clinical responders and non-responders. Specifically, the ability to induce ATM activation (i.e. pATM) in CD8 T cells during the proliferative response to PD-1 blockade was substantially more robust in clinical responders and nearly absent in most clinical non-responders. Together these data not only establish a platform with which to assess the role of DDR in human T cells, but they also point to the possible clinical utility of monitoring genome integrity surveillance in immune cells during immunotherapy in cancer patients.

The disconnect between immunological and clinical response in this uterine cancer cohort prompted a deeper examination the molecular pathways in CD8 T cells responding to PD-1 blockade. Recent studies have highlighted the importance of genome integrity maintenance in CD8 T cells responding to viral infection in mice and have identified distinct DNA damage circuits in different subsets of CD8 T cells (20). How cell division induced by checkpoint blockade such as anti-PD-1 is connected to CD8 T cell-intrinsic DDR pathways has not been previously examined in detail. This topic perhaps has particular relevance in cancer where many patients are treated with chemotherapy or other regimens that may impact genomic integrity, not only of cancer cells, but also of immune cells. In addition, many T cell types relevant for immunotherapy including T_EX_ have already undergone extensive activation and previous cell division, the consequences of which for DDR are unclear. By combining high resolution analysis of CD8 T cell subsets and differentiation states with protein markers of response to different types of DNA damage induction, we revealed different patterns of genomic integrity surveillance or damage in different subtypes of CD8 T cells. It will be important in the future to further interrogate the temporal kinetics of such responses and link these DDR signatures to patterns of patient pretreatment in larger cohorts. Moreover, the ability to monitor DDR in subtypes of CD8 T cells might provide insights about the fitness of responding T cell populations upon stress and suggest additional treatment modalities or inform combination therapies. For example, these data may have implications for combinations of genotoxic therapies such as chemotherapy and radiotherapy with checkpoint blockade. Our data suggest that signatures of DDR in immune cells in such settings might be important indicators of which patients will benefit.

As a small number of patients in the MSI-H cohort examined here had gMMRd. Although the number of these patients in the current cohort was too small for robust statistical analysis, it is possible that germline DNA repair deficiencies could have distinct impacts on the ability of T cells to manage genomic integrity. It is clear that gMMRd do not have overt profound immune deficiencies. These patients have no obvious clinical manifestations of poor responses to infectious agents of vaccine induced immunity. However, it is possible that the chronic nature of the response to a tumor adds additional stress on the DDR in tumor specific T cells leading to a role for DDR in immunotherapy that is distinct in gMMRd patients compared to non-gMMRd patients. This issue may be further complicated by the different genes that constitute gMMRd (e.g. *MSH2, MLH1, MSH6, EPCAM* etc.) and greater or lesser roles for these genes in T cells compared to tumor cells. Nevertheless, analysis in larger cohorts is warranted to further dissect the impact of gMMRd on differentiation and function of immune cells including T cells.

The identification of pATM as a DDR signature of relevance in the setting of PD-1 blockade in hypermutated uterine cancer patients may provide additional mechanistic insights. For example, induction of pATM in Ki67+ CD8 T cells at the peak of the response to PD-1 blockade distinguished clinical responders and non-responders. Indeed, the transcriptional program of proliferating CD8 T cells in the blood of MSI-H uterine cancer patients who benefited from anti-PD-1 treatment was enriched in an ATM-dependent gene expression signature. One obvious interpretation of these data is the connection between pATM, and repair of DNA damage lesions associated with the proliferative response. Such DDR may enable sustained CD8 T cell responses and more efficient anti-tumor activity whereas an inefficient repair process would be associated with induction of senescence and/or death of the responding T cells. There are also other possibilities. For example, although ATM has a well-described role in DNA repair, ATM signaling can be induced directly by TCR stimulation (39, 56). Whether this pATM induction is only due to cell cycle associated DNA damage downstream of TCR signaling or via other TCR signaling pathways is unclear. Moreover ATM signaling can regulate the Akt-mTOR pathway (57), and ATM can also be activated without DSB by oxidative stress (58). Thus, ATM may have a key role in managing genomic DNA damage, but this response might also reflect additional features of T cell fitness under stress. Nevertheless, these data highlight the DDR axis and a connection to ATM in different CD8 T cell subsets in response to PD-1 blockade immunotherapy in uterine cancer patients.

There are several potential clinical implications of these findings. First, with further examination, it may be possible to test the utility of an immune cell-intrinsic DDR-based biomarker for early on-treatment identification of patients likely to experience clinical benefit or not. Second, these data may have implications for testing future immune checkpoint combinations, including combinations with DNA damaging cancer treatments such as chemotherapy, targeted inhibitors such as PARP inhibitors or radiation. It may be informative to examine timing and sequencing of such genotoxic therapies used with immunotherapy in future studies. Applying the DDR profiling approach developed here may provide insights into how to best achieve anti-tumor effects while preserving immune cell fitness. Finally, it will be valuable to examine whether these results can extend to other cancers where the immunological response to PD-1 blockade also does not perfectly correlate with clinical response.

This study has several limitations. First, the cohort comprises a relatively small sample size, and high-quality samples were not available at all time points for all assays, thus limiting statistical power in some cases. Second, although the flow cytometric DDR panel allow insights into several DDR pathways in T cells, we were limited in the number of proteins and pathways that could be examined with this platform and traditional biochemistry is not possible because of small numbers of proliferating CD8 T cells available. Third, these data are limited to peripheral blood. Despite the benefit and ease of detecting pharmacodynamic responses to PD-1 blockade in the blood(7-9,59), it will be important in the future to examine these types of responses in the tumor microenvironment and correlate with what is occurring in the periphery. Finally, we recognize that there are other immune cell types that may be important contributors to immunological and clinical response to PD-1 pathway blockade that were not examined here. Overall, this study is hypothesis-generating, and will benefit from further examination in a larger cohort, including sufficient numbers of patients with germline DDR defects.

In summary, with the development of DDR-Immune platform, we identify CD8 T cell-intrinsic DDR as a possible previously unappreciated independent determinant of clinical response to immunotherapy. These observations provide insights into the disconnect between immunological and clinical response in highly mutated cancers. Moreover, the identification of a key role for DDR responses in immune cells and association with outcomes has implications for the relationships between genotoxic cancer treatments and immunotherapies. Finally, these studies also highlight the possibility that targeting immune cell DDR to improve responses to genomic damage could have therapeutic benefit.

## ACNOWLEDGEMENTS

We thank the patients and donors. We thank the members of the Wherry laboratory for helpful discussion. We acknowledge the Penn CMB microscopy core (Xinyu Zhao) for imaging immunofluorescence. We thank Clara R. Malekshahi and Daniel P. Beiting for help with RNAseq. We thank Penn Human Immunology Core (HIC) for healthy donor PBMC. We would like to thank the patients and their families for their time and effort in participating in the clinical trial, as well as the MSKCC clinical coordinators and team for helping to summarize clinical information and preparing clinical specimens.

## AUTHORS CONTRIBUTIONS

YM, DZ, CCF and EJW conceptualized the study and designed experiments. YM developed DDR-Immune platform. YM performed experiments with support from LC, NW and DM. YM analyzed the data with support from SM and DAO. NW generated CRISPR knock-out human T cells for ATM gene with support from CHJ. CX and YM performed immunofluorescence (IF) with support from SLB. ACH and RSH provided intellectual input. CCF and DZ designed and implemented clinical trial at MSKCC (NCT03241745). ARG edited the manuscript. YM, DZ, CCF and EJW wrote and edited the manuscript. All authors reviewed and accepted the final manuscript.

## MATERIALS AND METHODS

### Healthy Donor PBMC

Blood was acquired with informed consent of all study participants and with the approval of the University of Pennsylvania Institutional Review Board. Healthy adult human peripheral blood mononuclear cells were obtained from the Human Immunology Core (HIC) of University of Pennsylvania. Peripheral blood mononuclear cells (PBMCs) were isolated using Ficoll (GE Healthcare) gradient and stored using standard protocols.

### Patients and specimen collection

Patients with MSI-H, MMRd or hypermutated uterine cancers were enrolled for treatment with nivolumab (480 mg/kg by infusion every 4 weeks) on NCT03241745 at Memorial Sloan Kettering Cancer Center (MSKCC). Patients consented for blood collection under protocol IRB#17-180 at MSKCC, in accordance with the Institutional Review Board of the institution. The clinical follow-up data was last updated in January 2022. Peripheral blood was obtained in sodium heparin tubes before treatment and before each nivolumab infusion every 4 weeks for 12 weeks. Peripheral blood mononuclear cells (PBMCs) were isolated using Ficoll gradient and stored using standard protocols.

### DNA-damage treatments

T cells were isolated using EasySep™ Human T Cell Isolation Kit (Stem Cell) as per manufacturer’s instructions, from single cell suspensions of human PBMC, prior to the following DNA-damage treatments.

#### *Ex vivo* irradiation (“IR”)

Single cell suspensions of human PBMC were exposed to 0 or 500 rad (5 Gray (Gy)) of gamma irradiation by Gammacell® 40 Exactor (MDS Nordion).

#### *Ex vivo* UV irradiation (“UV”)

Single cell suspensions of human PBMC were exposed to 3000 μw/cm2 (= 3 J/m2) UV light (UVB: 312 nm) by FOTO/UV21 Transilluminators (FOTODYNE incorporated) for 30 minutes.

#### Anti-CD3/CD28 stimulation (“aCD3/CD28”)

Single cell suspensions of human PBMC were stimulated with Dynabeads™ Human T-Activator CD3/CD28 for T Cell Expansion and Activation (ThermoFisher Scientific) as per manufacturer’s instructions.

After the treatments, cells were incubated at 37°C in RPMI supplemented with 10% FBS, 2 mM L-Glutamine, 100 U/ml Penicillin, and 100 U/ml Streptomycin. Cells were then harvested at the indicated time points (30 minutes, 2, 4, 24, 48, 72 hours post-treatment). When not specified, cells were collected at 2 hours post-irradiation (IR), 4 hours post-UV-irradiation, and 24 hours post-aCD3/CD28 stimulation.

### Flow cytometry

PBMCs were stained with a master mix of antibodies listed in **Supplementary Table 5**. Permeabilization was performed using the Foxp3 Fixation/Permeabilization Concentrate and Diluent kit (eBioscience). Cells were resuspended in 1% para-formaldehyde until acquisition on a BD Biosciences Symphony cytometer and analyzed using FlowJo (BD Biosciences). Caspase 3 staining was performed using PE Active caspase 3 Apoptosis Kit (BD) as per manufacturer’s instructions, except the intracellular staining was performed for 1 hour in 4 °C (60).

### RNA extraction, cDNA preparation and qPCR

CD8 T cells were isolated with EasySep™ Human CD8+ T Cell Isolation Kit (Stem Cell) and treated with DNA-damage treatments described above. After the treatment, T cells were collected, and RNA extraction was performed using RNeasy Plus Micro Kit (50) (Qiagen) as per manufacturer’s instructions. Isolated RNA was quantified with NanoDrop (ThermoFisher), and reverse transcribed to cDNA with RNA to cDNA EcoDry Premix (Random Hexamers) (Takara/Clonetech). qPCR mastermix was made with Eukaryotic 18S rRNA Endogenous Control (ThermoFisher), Universal Master Mix (Taqman) (ThermoFisher), and the following primers (ThermoFisher). Parp1 (Hs00242302_m1), Msh2 (Hs00953527_m1). qPCR was performed with QuantStudio 6 Flex Real-Time PCR System (ThermoFisher).

### Immunofluorescence (IF)

Cells were fixed in 4% paraformaldehyde in PBS for 30 minutes at room temperature. After washing twice in PBS, cells were permeabilized with 0.5% Triton X-100 in PBS for 10 minutes, followed with two PBS washes. Cells were then blocked in 10% BSA in PBS for 1 hour at room temperature, and were incubated with γH2AX antibodies (Abcam, ab2893, 1:200 or Millipore, 05-636, 1:100) and anti-ATM (phospho S1981) antibody (Abcam, ab81292) in 5% BSA in PBS supplemented with 0.1% Tween 20 (PBST) for overnight incubation at 4℃. The next day, cells were washed 4 times for 10 minutes each with PBST, and then incubated with Alexa Fluor (AF)-conjugated secondary antibodies (AF488 and AF555) (Life Technologies) in 5% BSA/PBST for 1 hour at room temperature, followed by 4 times of 10-minute washes with PBST. Cells were then stained with 1 μg/ml DAPI for 5 minutes and washed twice with PBS. The coverslips were mounted with ProLong Gold (Life Technologies) and imaged with Leica TCS SP8 fluorescent confocal microscope (63X).

### Generation of gene-disrupted ATM-deficient T cells

#### Editing reagents and guide RNA design

Guide RNAs (sgRNAs) were designed using CHOPCHOP and the Broad Institute’s sgRNA design tool (doi.org/10.1093/nar/gkz365, https://portals.broadinstitute.org/gpp/public/analysis-tools/sgrna-design). Guide RNAs were picked based on their predicted efficiency, off-target reactivity, and location within the transcript. Chemically modified sgRNA’s (IDT) were screened for editing efficiency in primary human T cells as described below. The lead sgRNA used in experiments (5’TGATAGAGCTACAGAACGAAAGG’3) binds to exon 3 in the ATM transcript. A guide RNA targeting the irrelevant gene (EMX1) was used as a control non-targeting sgRNA as previously described(61). HiFi SpCas9 protein (SpyFi^TM^) was purchased commercially (Aldevron) and aliquoted to avoid frequent freeze/thawing.

#### T cell isolation and culture

Human T cells were procured through the University of Pennsylvania Human Immunology Core. Primary human CD4 and CD8 T cells were isolated from PBMC of healthy donors using commercially available CD4 and CD8 selection kits (Miltenyi). CD4 and CD8 T cells were cultured at 1×10^6^ cells/ml at a 1:1 CD4/CD8 ratio in R10 (RPMI, 10% FBS, 1% HEPES, 1% GlutaMAX, 1% Penicilin/Streptomycin) supplemented with 5 ng/ml human IL-7 and human IL-15 (Peprotech) each. After nucleofection, T cell were activated for 120 hours with anti-CD3/anti-CD28 dynabeads (ThermoFisher) at a 3:1 bead/cell ratio. After 120 hours, dynabeads were magnetically removed and T cells were cultured in standard culture flasks at 1×10^6^ cell/ml in R10 supplemented with IL-7 and IL-15 until growth kinetics and cell size indicated that cells have rested from stimulation (∼350 fL).

#### T cell nucleofection

T cell nucleofection was carried out as previously described (62). 5-10×10^6^ non-activated T cells were washed in serum-free minimal essential media (ThermoFisher) and resuspended in 100 μL P3 nucleofection solution (Lonza). T cells were then electroporated with ribonucleoprotein (RNP) complexes of sgRNA and SpCas9 in 100 μL cuvettes using pulse code EH111 of the 4D nucleofector (Lonza). RNP complexes were generated by incubating 5 ug of chemically modified sgRNA with 10 ug HiFi SpCas9 for 15 minutes at room temperature. After nucleofection, T cells were allowed to recover for 48 hours in R10 supplemented with IL-7 and IL-15 at 30 °C. After recovery, T cell were activated and expanded as described above.

#### Genomic DNA analysis

At the end of T cell expansion, genomic DNA was isolated from 3×10^6^ cells by spin-column based DNA purification according to the manufacturer instruction (DNeasy, Qiagen). Knockout efficiency was analyzed on the genomic level through TIDE analysis (Tracking of Indels by DEcomposition). INDEL frequency was measured on the genomic level by PCR amplification (500-1000 bp) of the CRISPR-targeted loci (Fwd: 5’ GCTACTACTGCAAGCAAGGCA’3, Rev: 5’AAATGCCAAATTCATATGCAAGGC3’). DNA was amplified using AccuPrime Taq polymerase (Invitrogen). PCR amplicons were purified from 1% agarose gels following electrophoresis by column purification (Takara) and submitted for sanger sequencing through Genewiz (Primer: 5’ GCCTTTGACCAGAATGTGCC’3). Sanger sequencing trace files of control and gene-edited PCR amplicons were uploaded to the web app TIDE (Tracking of Indels by DEcomposition) for analysis(63).

#### Western blotting

For western blotting, 5×10^6^ T cells at the end of expansion were harvested from culture and washed in cold PBS (ThermoFisher). The cell pellet was resuspended in ice-cold 1x RIPA buffer (Cell Signaling Technologies) supplemented with protease and phosphatase inhibitor cocktail (ThermoFisher). The lysates were incubated on ice while being vortexed every 10 minutes for 5 seconds intervals. The lysates were then centrifuged at 16,000 g for 15 minutes in a refrigerated centrifuge. The supernatants were collected, and protein concentration was measured by DC protein assay according to the manufacturer’s instructions (BioRad). NuPAGE LDS sample buffer and reducing agent (Invitrogen) were added to equal amounts of protein lysates and boiled at 95 °C for 10 minutes. Proteins were separated using NuPAGE 4-12% Bis-Tris gels in the XCell SureLock electrophoresis chambers for 2 hours at 100 V. After electrophoresis, proteins were transferred to a methanol-activated PVDF-FL membrane (Millipore Sigma) in NuPAGE transfer buffer (Invitrogen) for 2 hours at 4 °C using Mini-trans blot wet tank transfer (BioRad). After transfer, membranes were blocked with TBS blocking buffer (Licor) for 30 minutes at room temperature. Proteins were visualized by incubating membranes with primary antibodies against ATM (Clone D2E2, 1:1000, CST), pATM-Ser1981 (Clone D25E5, 1:1000, CST), and beta-actin (Clone 8H10D10, 1:2000, CST), diluted in TBS antibody diluent (Licor) and incubated overnight at 4 °C. Each membrane was only incubated with one primary antibody from a single origin species (rabbit or mouse) at one time. The membranes were washed in TBS-T (Cell Signaling Technologies) washing buffer and incubated with donkey anti-rabbit or donkey anti-mouse secondary antibodies (1:10,000; IRDye 680RD or IRDye 800CW, LI-COR) for 45 minutes at room temperature. The membranes were then washed again in TBS-T and imaged with an Odyssey CLx (LI-COR). Fluorescence was evaluated with Image Studio (LI-COR). When membranes were reused to visualize additional proteins, membranes were incubated in stripping buffer (ThermoFisher) for 30 minutes, washed in TBS-T and re-blocked in TBS Odyssey blocking buffer (LI-COR) for 1 hour at room temperature and then processed as described above.

### RNAseq sample preparation and sequencing

CD3 T cells were enriched using EasySep™ Human T Cell Isolation Kit (Stem Cell) as per manufacturer’s instructions, prior to sorting CD8 T cells from ATM^WT^ and ATM^KO^ T cells. RNA was extracted in RLT buffer supplemented with beta-mercaptoethanol and processed with the RNeasy Micro Kit (QIAGEN) as per the manufacturer’s instructions. Total RNA libraries were prepared using the SMARTer Stranded Total Kit v3 (Takara). Extracted RNA and libraries were assessed for quality on a TapeStation 2200 instrument (Agilent). RNA libraries were quantified using the KAPA Library Quantification Kit and sequenced on an Illumina NextSeq 500 instrument (150 base pairs, paired-end) on mid-output flow cells.

### RNAseq data processing and analysis

FASTQ files were aligned using STAR 2.5.2a against the hg38 human reference genome. The aligned files were processed using PORT gene-based normalization (https://github.com/itmat/Normalization). Differential gene expression was performed with Limma. *removeBatchEffect* function was used to fit the linear model for the RNAseq expression data to remove unwanted batch effect from sample batches. Limma-voom was used to identify transcripts that were significantly differentially expressed between experimental groups using an adjusted p-value <0.05.

### Single-cell RNAseq sample preparation and sequencing

PBMC samples from patients with MSI-H, MMRd or hypermutated uterine cancers, collected before therapy and 2 weeks after anti-PD-1 blockade therapy initiation, were processed using EasySep™ Human T Cell Isolation Kit (Stem Cell) as per manufacturer’s instructions. Non-naive CD3 T cells were sorted into 1.5 ml Lo-Bind Eppendorf tubes with complete RPMI (10% FBS) and were washed with PBS twice before loading to a Chromium single cell sorting system (10× Genomics). Library construction was performed following the protocol of Chromium Next GEM Single Cell 5’ Kit v2, with a standard loading targeting 1 × 10^4^ cells recovered. The final pooled library with 16 samples was sequenced on NovaSeq 6000 instrument (100 cycles, paired-end) on S2 v1.5 flowcells.

### Single-cell RNAseq data processing and analysis

BCL files from sequences samples were demultiplexed, aligned to the human hg38 genome, filtered, and UMI counted using Cell Ranger Software v4.0.0 (10x Genomics). Downstream analysis was performed using Seurat v4.0.1(64). Briefly, individual library outputs from the Cell Ranger pipeline were loaded, filtered to remove cells with > 5% mitochondrial reads. Filtered data from individual libraries were imputed using SAVER v0.3.0 (65) with default settings. Imputed count matrices were used for subsequent analysis in Seurat. Single-cell enrichment scores were calculated using the *AddModuleScore* function in the Seurat package. Cell cycle scores were calculated using the *CellCycleScoring* function in the Seurat package using previously defined gene signatures for each cell cycle stage (66). Differential gene expression was conducted using the *FindMarkers* function in Seurat package and a gene signature was created using a threshold for adjusted p-value at 0.05 and log2(fold change) >0.25. Top differential genes by significance for each cluster were visualized as heatmap of scaled expression using DoHeatmap function from Seurat package. Expression of various genes were projected on to the UMAP along with other clinical information. Percent composition of cells of various clinical groups were visualized using barplots from DittoSeq R package.

### Geneset enrichment analysis

GSEA(67) was performed using the Broad Institute software GSEA 4.0.3 (https://www.broadinstitute.org/gsea/index.jsp). Enrichment scores were calculated by using a ranked list of genes for a pairwise comparison using the gene signature created from Single cell RNAseq analysis as described above.

### TCGA dataset, Gene Set Variation Analysis, survival analysis and sample size calculations

TCGA dataset of UCEC was downloaded from Broad Institute website (http://gdac.broadinstitute.org/). *getTCGA* function from R package *TCGA2STAT_1.2* was used to get UCEC, COAD and SKCM RNAseq2 expression data. For MSI status determination, *GDCprepare_clinic (query, “msi”)* function of *TCGAbiolinks_2.16.4* package was used, then merged with RNAseq2 expression data for further analysis. For Gene Set Variation Analysis (GSVA) analysis, *GSVA_1.36.2* R package was used(54). For survival analysis, *survival_3.2-7* (https://cran.r-project.org/package=survival) and *survminer_0.4.*9 (https://cran.r-project.org/package=survminer) functions in R were used. The optimal cutpoint was calculated by *cutoffr_1.1.1* (https://cran.r-project.org/package=cutpointr). Nominal significance for survival analysis was calculated by log-rank test, and the adjustment for multiple comparisons was performed with Bonferroni correction. Sample size analysis for Log_2_(FC pATM) was performed using the online UCSF Sample Size Calculator (https://sample-size.net/sample-size-survival-analysis/) with parameters α=0.05 (corresponding to a nominal significance threshold P < 0.05), β=0.20 (corresponding to 80% power), q1=0.4666, and HR=2. Cox proportional hazard modeling to estimate the hazard ratio (HR) was calculated by *coxph* function in R (https://www.rdocumentation.org/packages/survival/versions/3.2-13/topics/coxph) after determining the optimal cutpoint for Log_2_(FC pATM). R versions 4.0.4 and 3.5.3 were used.

### Statistical analysis

Mann-Whitney tests (paired and unpaired), two-sided Wilcoxon matched-pairs test, ANOVA analyses with Dunnett’s test, one-sample t-test and paired t-test, and area under the curve (AUC) calculations were performed using Prism software (GraphPad Software Inc. of La Jolla, CA). Correlation analysis and Fisher exact test were performed as previously described(68), using the NonParametricCorrelationCode (https://github.com/wherrylab/statistics_code).

### DATA AVAILABILITY

10x scRNAseq of PBMC CD8 T cells samples from patients with MSI-H, MMRd or hypermutated uterine cancers, and bulk RNAseq of ATM^WT^ and ATM^KO^ T cells will be deposited to Gene Expression Omnibus (GSE) upon publication.

## Supplementary Figure legends

**Supplementary Fig. 1: Comparison of clinical features between clinical responders and non-responders.**

**(A-C)** Comparison of TMB **(A)**, age **(B)** and days since last chemotherapy treatment to the initiation of nivolumab **(C)**, between responders and non-responders (Responders (R: blue): N = 20, Non-Responders (NR: red): N = 14). Clinical response was determined by the best response by RECIST criteria. For a violin plots, dashed line denotes median and dotted lines denote quantiles. p-values determined by unpaired Mann-Whitney U-test.

**Supplementary Fig. 2: CD8 T cell proliferative kinetics of healthy donors.**

Fold change of Ki67 by CD8 T cells over 3 weeks in healthy donors (n = 7) as described (9). Error bars denote SD, center line denotes mean.

**Supplementary Fig. 3: scRNAseq of CD8 T cells from PBMC of MSI-H endometrial cancer patients.**

**(A, B)** UMAP projection of CD8 T cell scRNA-seq data, colored by time points pre or 2 weeks post PD-1 blockade (A), and clinical response by RECIST criteria (B). **(C)** UMAP projection of CD8 T cell scRNA-seq data, colored by expression of representative genes. **(D)** Violin plots depicting expression of representative genes within Louvain clusters. **(E)** Line plot of the frequency of cluster 2 per patient, pre- and post PD-1 blockade. (N = 7, 4 responders (1 responder did not have cells in cluster 2), 3 non-responders). p-values determined by Wilcoxon matched test.

**Supplementary Fig. 4: Gating strategies for human CD8 T cell subsets.**

Gating strategies for CD8 T cells, gated on live CD3+CD8+ T cells. Healthy donor PBMC.

**Supplementary Fig. 5: Fold change of DDR markers in different CD8 T cell subsets compared with naive CD8 T cells after DNA damage treatment.**

**(A, B)** 2 hours post IR. Fold change increase of γH2AX **(A)** and pATM **(B)** compared with naive CD8 T cells. **(C, D)** 4 hours post UV. Fold change increase of γH2AX **(C)** and pATM **(D)** compared with naive CD8 T cells. **(E, F)** 24 hours post aCD3/CD28 stimulation. Fold change increase of PARP1 **(E)** and MSH2 **(F)** compared with naive CD8 T cells. Error bar denotes mean ± SD. * p<0.05, ** p<0.01, *** p<0.001, **** p<0.0001 by one-sample t test.

**Supplementary Fig. 6: CD8 T cell subsets have different sensitivity to cell death in T cells.**

CD8 T cells were treated with IR and incubated for 24 hours. Cell viability and activated caspase 3 expression were assessed by flow cytometry.

**(A)** Representative flow plots of untreated and irradiated CD8 T cell subsets with live/dead (L/D) and activated caspase 3 staining. Alive = L/D-caspase 3-, Early apoptosis = L/D-caspase 3+, Late apoptosis =L/D+ caspase 3+, Necrosis = L/D+ caspase 3-. **(B)** Heatmap of row-normalized frequency of cell death across CD8 T cell subsets and DNA damage treatments. **(C)** Percentage of apoptosis (% caspase 3+) across different subsets in untreated (left) and irradiated (right) CD8 T cells. **(D, E)** Ratio of Difference (Δ) between the untreated and irradiation-induced γH2AX **(D)** and pATM **(E)** MFI Area under curve (AUC) in CD8 T cells through 24 hours post treatment, and % caspase 3 + of irradiated CD8 T cell subsets. (B-C) N = 5 per group, (C) Error bar denotes mean ± SD. * p <0.05, ** p <0.01, *** p<0.001 by one-way ANOVA with Dunnett’s test compared with naive CD8 T cells. (D, E) N = 3 per group, * p<0.05 by one-way ANOVA with Dunnett’s test compared with T_SCM_.

**Supplementary Fig. 7: Higher DDR marker expression in Ki67+ CD8 T cells.**

**(A)** Representative flow plot for γH2AX, PAPR and MSH2 expression at 2 weeks post initiation of nivolumab treatment; gated on Ki67+ CD8 (blue), Ki67-CD8 (green) and naive CD8 (gray). **(B)** Normalized MFI (nMFI) of the indicated markers in Ki67+ (blue) and Ki67− (green) CD8 T cells at 2 weeks post initiation of nivolumab treatment (N = 30). MFI values were normalized to those of naive CD8 T cells of healthy donor control. For γH2AX, %high γH2AX is also shown. **(C)** Representative flow plot for pATM, γH2AX, PARP1 and MSH2 expression at baseline prior to initiation of nivolumab treatment; gated on Ki67+ CD8, Ki67-CD8 and Naive CD8 T cells. **(D)** nMFI of the indicated markers in Ki67+ (blue) and Ki67− (green) CD8 T cells at baseline prior to initiation of nivolumab treatment (N = 31). For γH2AX, % high γH2AX is also shown. ** p<0.01, and *** p<0.001, **** p < 0.0001 by Wilcoxon matched-pairs signed rank test.

**Supplementary Fig. 8: Correlation matrix of clinical parameters, DDR markers and immune features in Ki67+ CD8 T cells at 2 weeks post initiation of nivolumab treatment.**

Immunological marker expression shown as % positivity of Ki67+ CD8 T cells. DDR features shown as normalized MFIs (nMFI), in which MFI values were normalized to those of naive CD8 T cells of healthy donor control. Spearman’s Rank Correlation coefficient (ρ) is indicated by square size and heat scale; significance indicated by: * p<0.05, ** p<0.01, and *** p<0.001; none of them reached FDR<0.05.

**Supplementary Fig. 9: Comparison of clinical features and CD8 T cell response between patients with germline MMR mutation and somatic mutations.**

**(A-C)** Comparison of TMB **(A)**, age **(B)** and days since last chemotherapy treatment to the initiation of nivolumab **(C)**, between patients with germline MMRd (gMMRd: orange) and without gMMRd (non-gMMRd: gray). **(D, E, F)** Ki67 expression in CD8 T cells at indicated times (gMMRd: N =5, non-gMMRd: N =28). **(G)** Comparison of fold change of Ki67 expression at peak of immunologic response versus pretreatment between gMMRd (orange) and non-gMMRd (gray). Dotted line denotes fold change of 1.73, which is the mean plus 2 standard deviations (SD) in healthy donors (see **Supplementary Fig. 2**) (gMMRd: N =5, non-gMMRd: N =27). **(H, I)** Representative flow plots **(H)** and percentages for T_EX_ (PD-1^hi^CD39^hi^ in non-naive CD8 T cells) **(I)** at baseline (Pre) and at 2 weeks post initiation of nivolumab treatment (Post) (gMMRd: N = 5, non-gMMRd: N =26). Error bar denotes mean ± SD. For violin plots (A-C, G, K, O), dashed lines denote median and dotted lines denote quartiles. * p <0.05, ** p <0.01, ***<0.001, **** < 0.0001 by unpaired Mann-Whitney U-test (A-C, G, I (Pre vs Pre, Post vs Post)), Wilcoxon matched-pairs signed rank test (E, F, I (Pre vs Post)).

**Supplementary Fig. 10: DDR marker expression kinetics in Ki67+ CD8 T cells between responders and non-responders.**

**(A-C)** Patterns of γH2AX **(A)**, PAPR1 **(B)** and MSH2 **(C)** expression kinetics in CD8 T cells in Responder versus Non-Responder (Responders (R: blue): N = 17, Non-Responders (NR: red): N = 13 (for MSH2: N =12)); P values were calculated using two-sided Wilcoxon matched-pairs test. * p <0.05. Error bar denotes mean ± SD. **(D-F)** Log fold change increase of γH2AX **(D)**, PARP1 **(E)** and MSH2 **(F)** from baseline to 2 weeks in Ki67+ CD8 T cells. (Responders (R: blue): n = 17, Non-Responders (NR: red): N = 13 (for MSH2 N = 12). P values were calculated using two-sided Wilcoxon matched-pairs test (A-C) and unpaired Mann-Whitney U-test (D-F). * p <0.05. Error bar denotes mean ± SD.

**Supplementary Fig. 11: Validation of ATM^KO^ primary human T cells by CRISPR.**

**(A)** ATM knock out (KO) does not impact primary human T-cell expansion *ex vivo*. Population doublings (left) and cell size (right) over the course of expansion. WT: wild type (transfected with gRNA of irrelevant gene), KO: knock out of ATM. **(B)** Verification of ATM knockout efficiency by sequencing, showing the frequency of insertion/deletion mutation induced by CRISPR. **(C-F)** Validation by Western blot. Relative signal intensity was normalized to beta actin control. **(C)** Representative Western blot image, and the quantification summaries for **(D)** relative signal intensity of ATM in non-irradiated T cells, **(E)** Ratio of relative signal intensity of phosphorylation of ATM (Ser 1981) (pATM) in non-irradiated ATM^KO^ T cells compared to ATM^WT^ T cells, and **(F)** relative signal intensity of phosphorylation of ATM (pATM) (Ser 1981) in irradiated ATM^WT^ T cells compared to non-irradiated ATM^WT^ T cells. Irradiated T cells (500 rad) were collected 1 hour post irradiation. **(G, H)** Validation by flow cytometry. **(G)** Representative flow cytometry plots for pATM expression in ATM^WT^ or ATM^KO^ CD8 T cells. Irradiated T cells (500 rad) were collected 1 hour post irradiation. **(H)** Delta MFI of pATM in irradiated CD8 T cells minus untreated T cells. * p<0.05, ** p<0.01, and **** p < 0.0001 by Wilcoxon matched-pairs signed rank test (A), one sample t-test (D-F), and paired t-test (H).

**Supplementary Fig. 12: Cell cycle analysis of scRNAseq from PBMC-derived CD8 T cells from hypermutated uterine cancer cohort.**

UMAP plot depicting cell cycle stage enrichment scores for each cell.

**Supplementary Fig. 13: Kaplan-Meier plots of the Overall Survival (OS) of MSI-H UCEC, COAD and SKCM TCGA dataset separated by ARGS score.**

**(A)** Kaplan-Meier plot of the Overall Survival (OS) between high (> 0.0023) versus low (< 0.0023) ATM-Responder Gene Signature (ARGS) score by Gene Set Variation Analysis (GSVA) of RNAseq2 expression data from MSI-H subset of The Cancer Genome Atlas Uterine Corpus Endometrial Carcinoma (TCGA-UCEC) data collection (N = 46). Blue line indicates ARGS score >0.0023, red line indicates ARGS score < 0.0023. Cut off was determined by *cutpointr* (see Methods). Nominal p-value was determined by log-rank test (p = 0.3, Bonferroni adjusted p= 1). **(B)** ARGS score comparison between patients with high *CD8A* expression (*CD8A* > median) versus low *CD8A*expression (*CD8A* < median) in TCGA-UCEC dataset. **(C)** Kaplan-Meier plot of the OS between high (>-0.1066) versus low (< −0.1066) ARGS score by GSVA of RNAseq2 expression data from TCGA Colon Adenocarcinoma (TCGA-COAD) data collection (N = 282). Blue line indicates ARGS score >-0.1066, red line indicates ARGS score < −0.1066. Cut off was determined by *cutpointr* (see Material and Methods). Nominal p-value was determined by log-rank test (p = 0.028, Bonferroni adjusted p= 1). **(D)** Kaplan-Meier plot of the OS between high (>-0.1269) versus low (< −0.1269) ARGS score by GSVA of RNAseq2 expression data from MSI-H subset of TCGA-COAD data collection (N = 49). Blue line indicates ARGS score >-0.1269, red line indicates ARGS score < −0.1269. Cut off was determined by *cutpointr* (see Methods). Nominal p-value was determined by log-rank test (p = 0.2, Bonferroni adjusted p= 1). **(E)** ARGS score comparison between patients with high *CD8A* expression (*CD8A* > median) versus low *CD8A* expression (*CD8A* < median) in TCGA-COAD dataset. **(F)** Kaplan-Meier plot of the OS between high (> 0.1468) versus low (< 0.1468) ARGS score by GSVA of RNAseq2 expression data from TCGA-SKCM data collection (N = 102). Blue line indicates ARGS score >0.1468, red line indicates ARGS score < 0.1468. Cut off was determined by *cutpointr* (see Methods). Nominal p-value was determined by log-rank test (p = 0.24, Bonferroni adjusted p= 1). **(G)** ARGS score comparison between patients with high *CD8A* expression (*CD8A* > median) versus low *CD8A* expression (*CD8A* < median) in TCGA-SKCM dataset. *p<0.05 by Wilcoxon matched-pairs signed rank test (B, E G)

## Supplementary Table Legends

**Supplementary Table 1: Clinical characteristics of MSI-H or hypermutated uterine cancer cohort.**

**Supplementary Table 2: Ranked gene lists of ATM^WT^ versus ATM^KO^ CD8 T cell genes.**

**Supplementary Table 3: Differentially Expressed Genes (DEG) from proliferating CD8 T cells of clinical responder versus non-responder patients.**

**Supplementary Table 4: Leading edge genes of GSEA analysis of ATM^WT^ versus ATM^KO^ CD8 T cells and DEG from proliferating CD8 T cells of clinical responder versus non-responder patients.**

**Supplementary Table 5: List of antibodies used for flow cytometry and DDR-Immune Platform.**

