## Supplementary Figures for "Induction of a CD8 T cell intrinsic DNA damage and repair response is associated with clinical response to PD-1 blockade in uterine cancer"

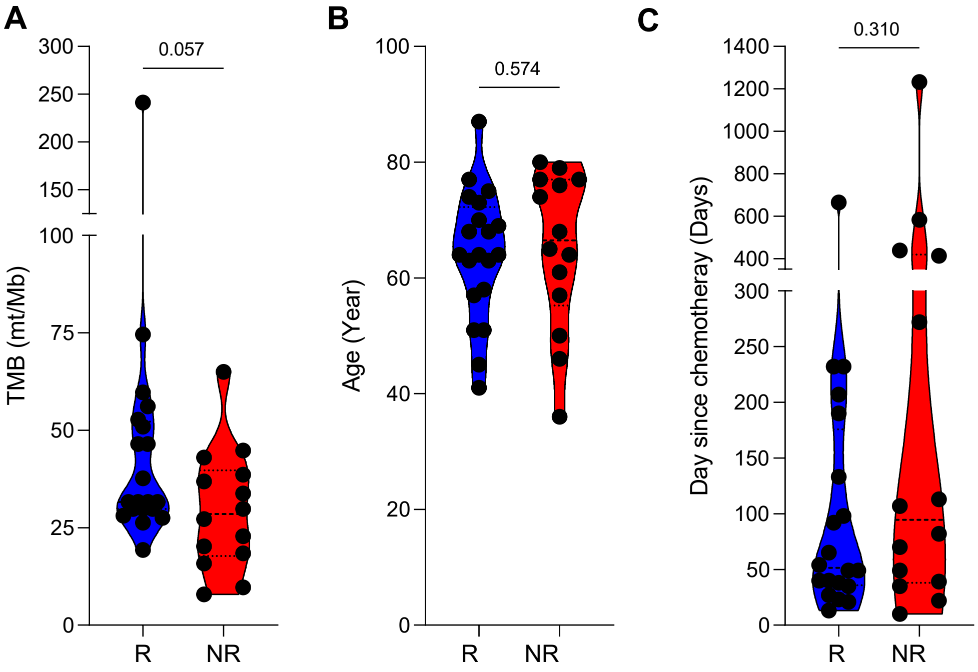
A
300
250
200
150
100
75
50
25
0
TMB (mt/Mb)
0.057
R
NR
B
100
80
60
40
20
0
Age (Year)
0.574
R
NR
C
1400
1200
1000
800
600
400
300
250
200
150
100
50
0
Day since chemotheray (Days)
0.310
R
NR

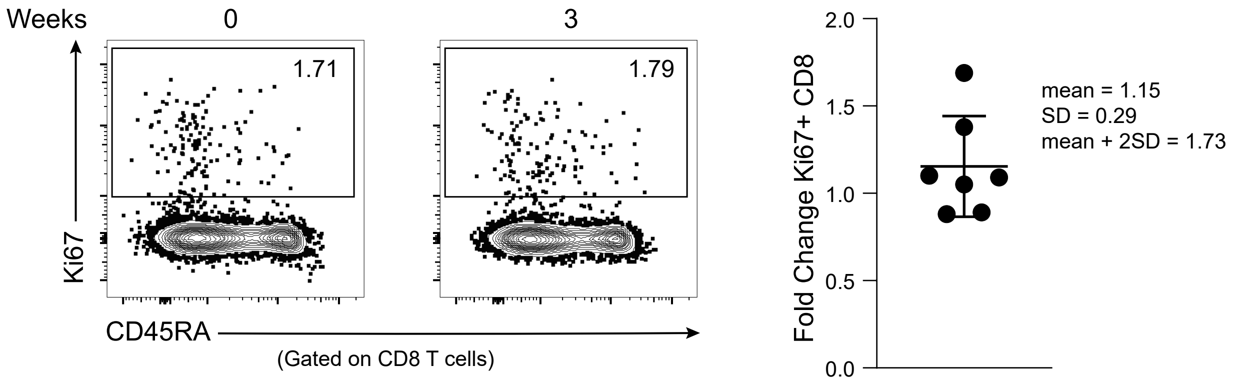
Weeks
0
3
1.71
1.79
Ki67
CD45RA
(Gated on CD8 T cells)
Fold Change Ki67+ CD8
2.0
1.5
1.0
0.5
0.0
mean = 1.15
SD = 0.29
mean + 2SD = 1.73

Supplementary Fig.3

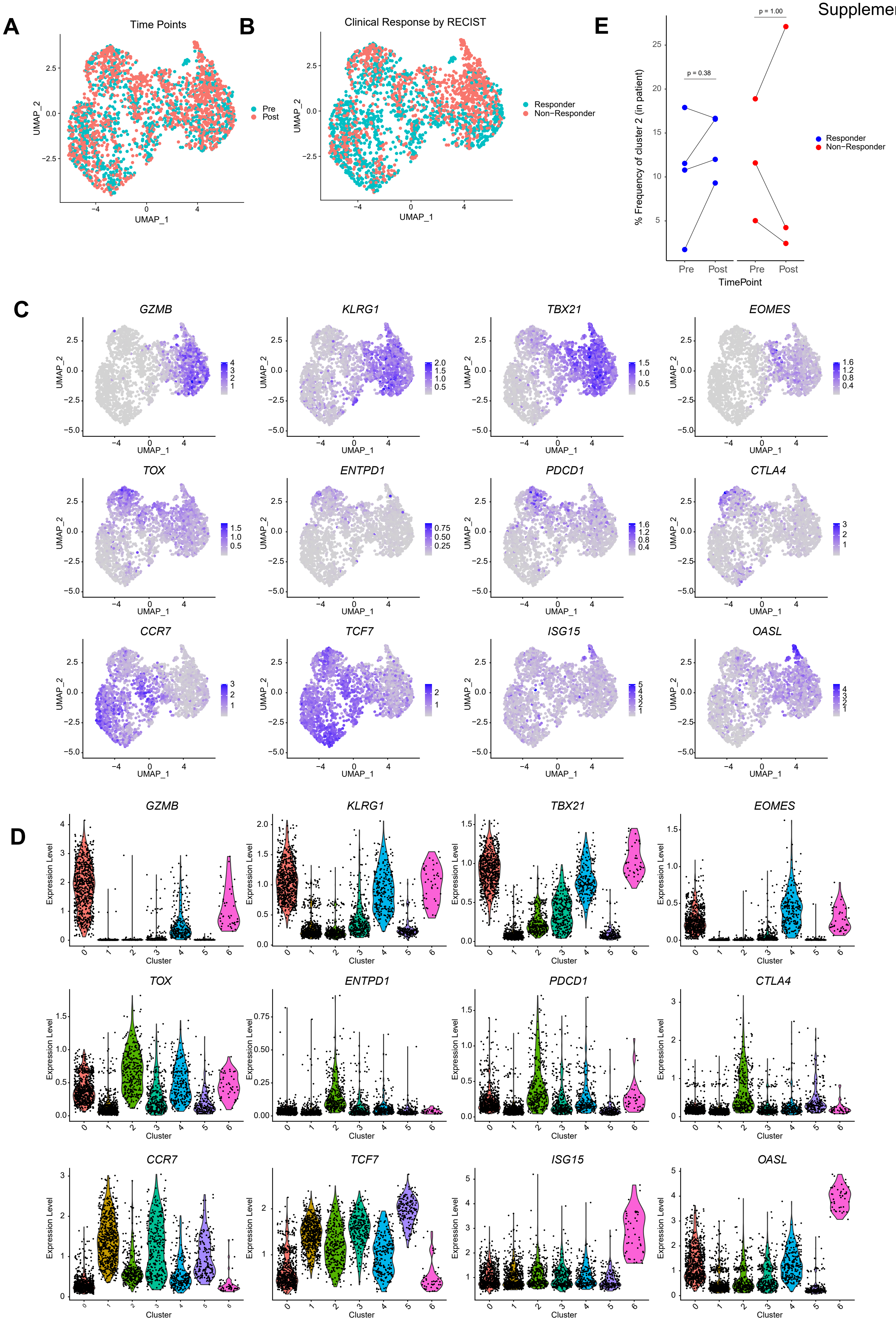

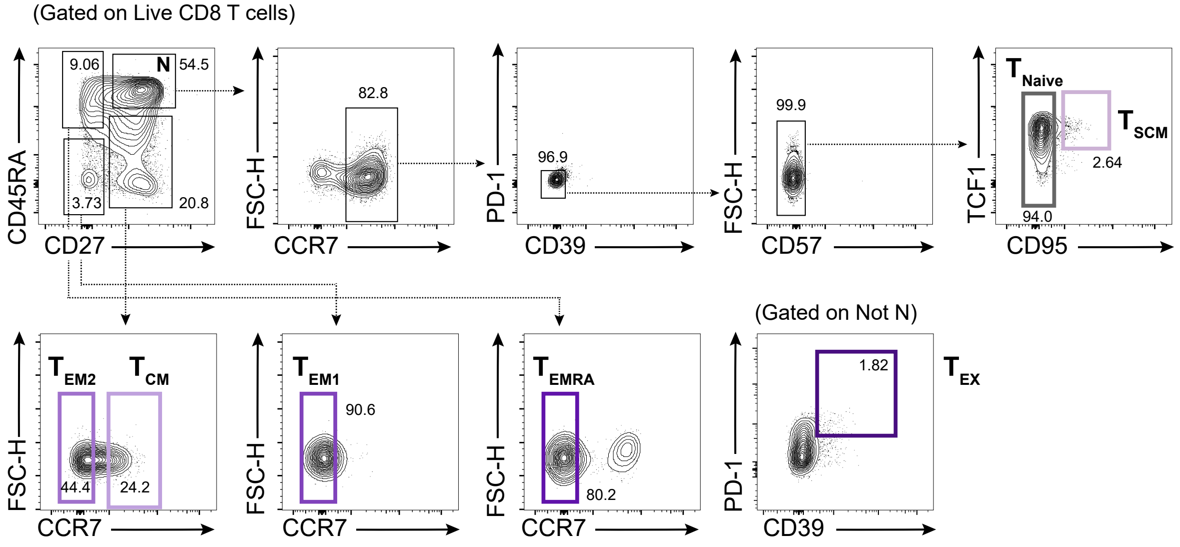
(Gated on Live CD8 T cells)
9.06
N
54.5
3.73
20.8
CD45RA
CD27
82.8
FSC-H
CCR7
96.9
PD-1
CD39
99.9
FSC-H
CD57
T Naive
T SCM
2.64
94.0
TCF1
CD95
T EM2
T CM
44.4
24.2
FSC-H
CCR7
T EM1
90.6
FSC-H
CCR7
T EMRA
80.2
FSC-H
CCR7
(Gated on Not N)
1.82
T EX
PD-1
CD39

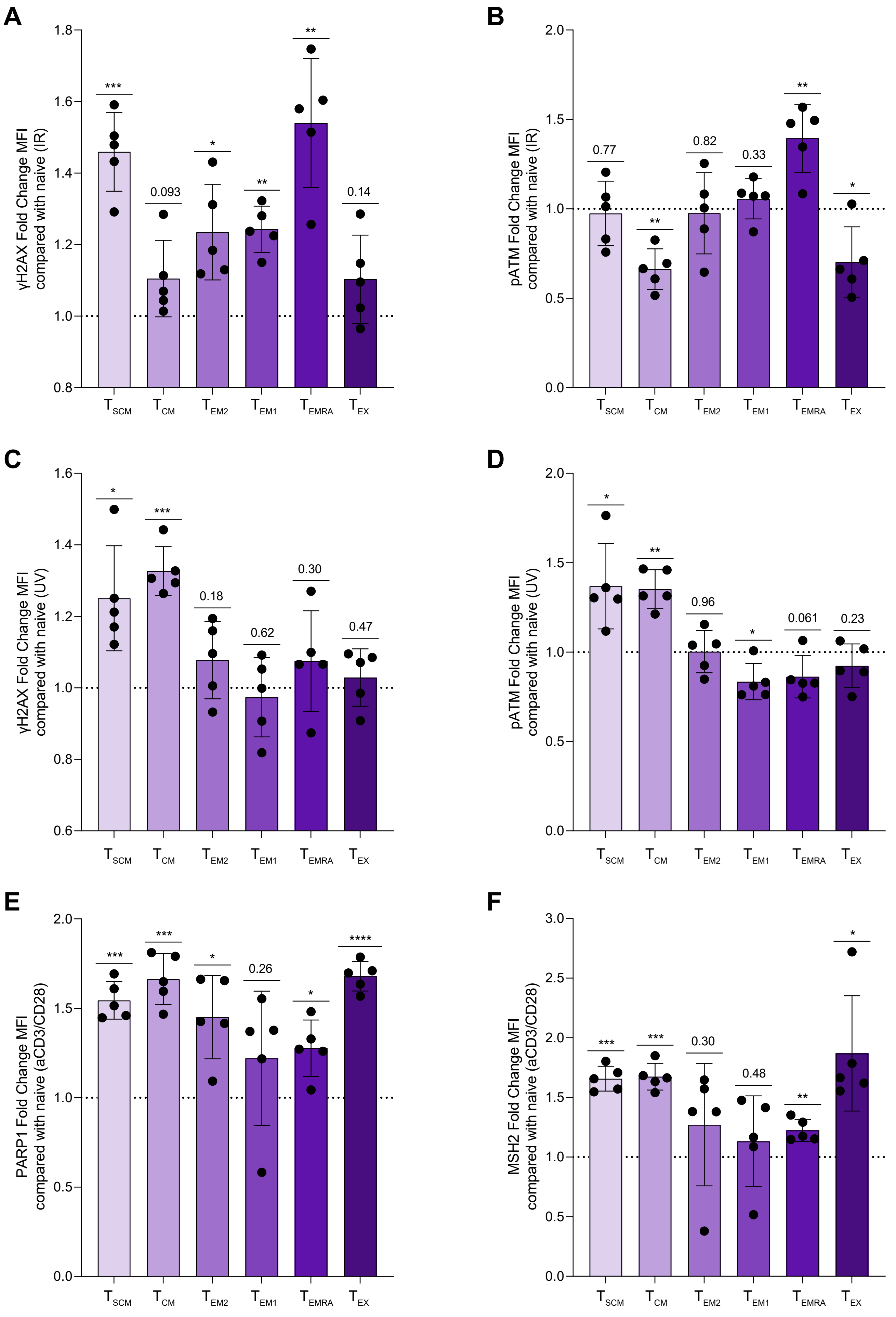
A
γH2AX Fold Change MFI compared with naive (IR)
1.8
1.6
1.4
1.2
1.0
0.8
***
0.093
*
**
**
0.14
T SCM
T CM
T EM2
T EM1
T EMRA
T EX
B
pATM Fold Change MFI compared with naive (IR)
2.0
1.5
1.0
0.5
0.0
0.77
**
0.82
0.33
**
*
C
γH2AX Fold Change MFI compared with naive (UV)
1.6
1.4
1.2
1.0
0.8
0.6
*
***
0.18
0.62
0.30
0.47
D
pATM Fold Change MFI compared with naive (UV)
*
**
0.96
*
0.061
0.23
E
PARP1 Fold Change MFI compared with naive (aCD3/CD28)
***
***
*
0.26
*
****
F
MSH2 Fold Change MFI compared with naive (aCD3/CD28)
3.0
2.5
2.0
1.5
1.0
0.5
0.0
***
***
0.30
0.48
**
*

Suplementary Fig.6

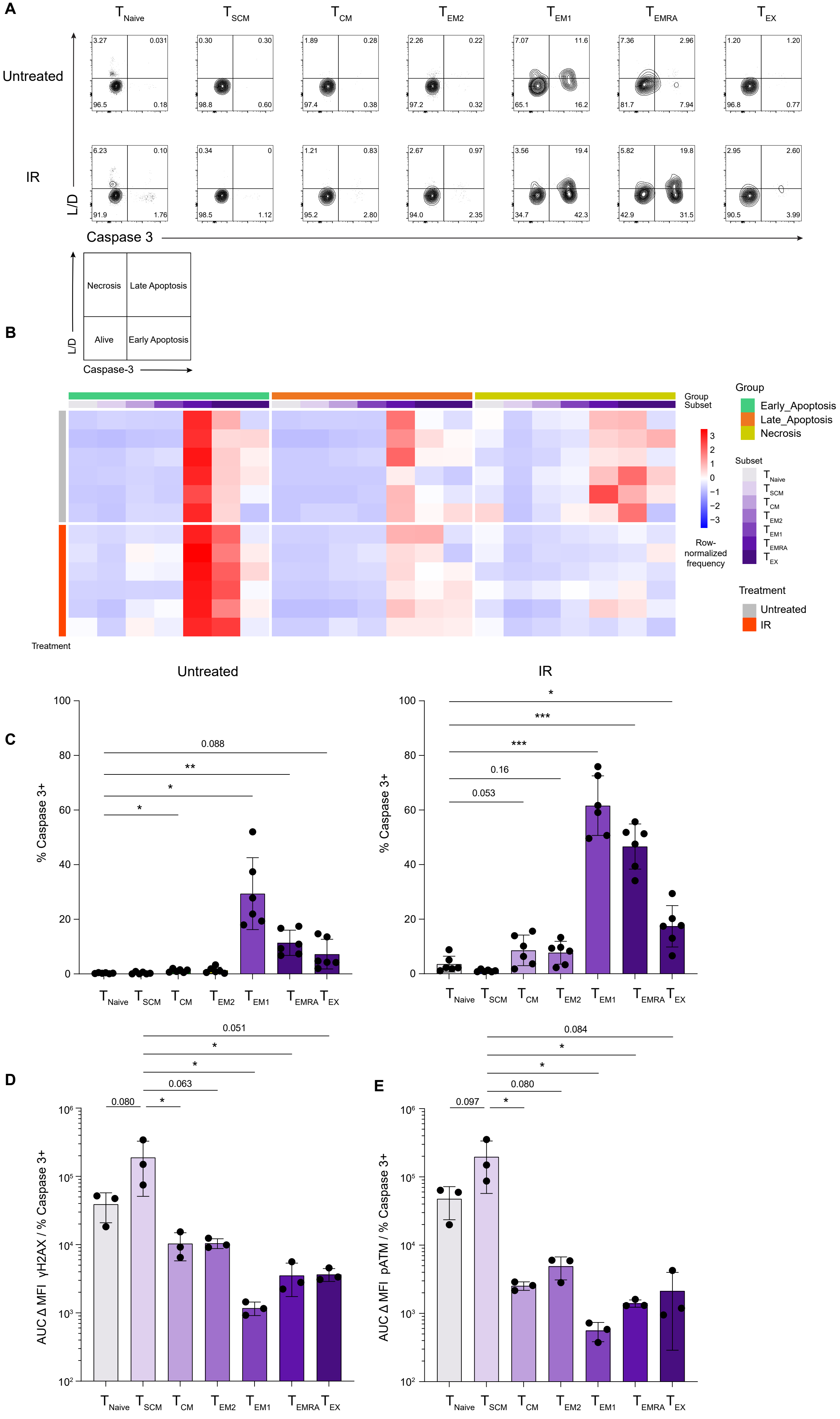

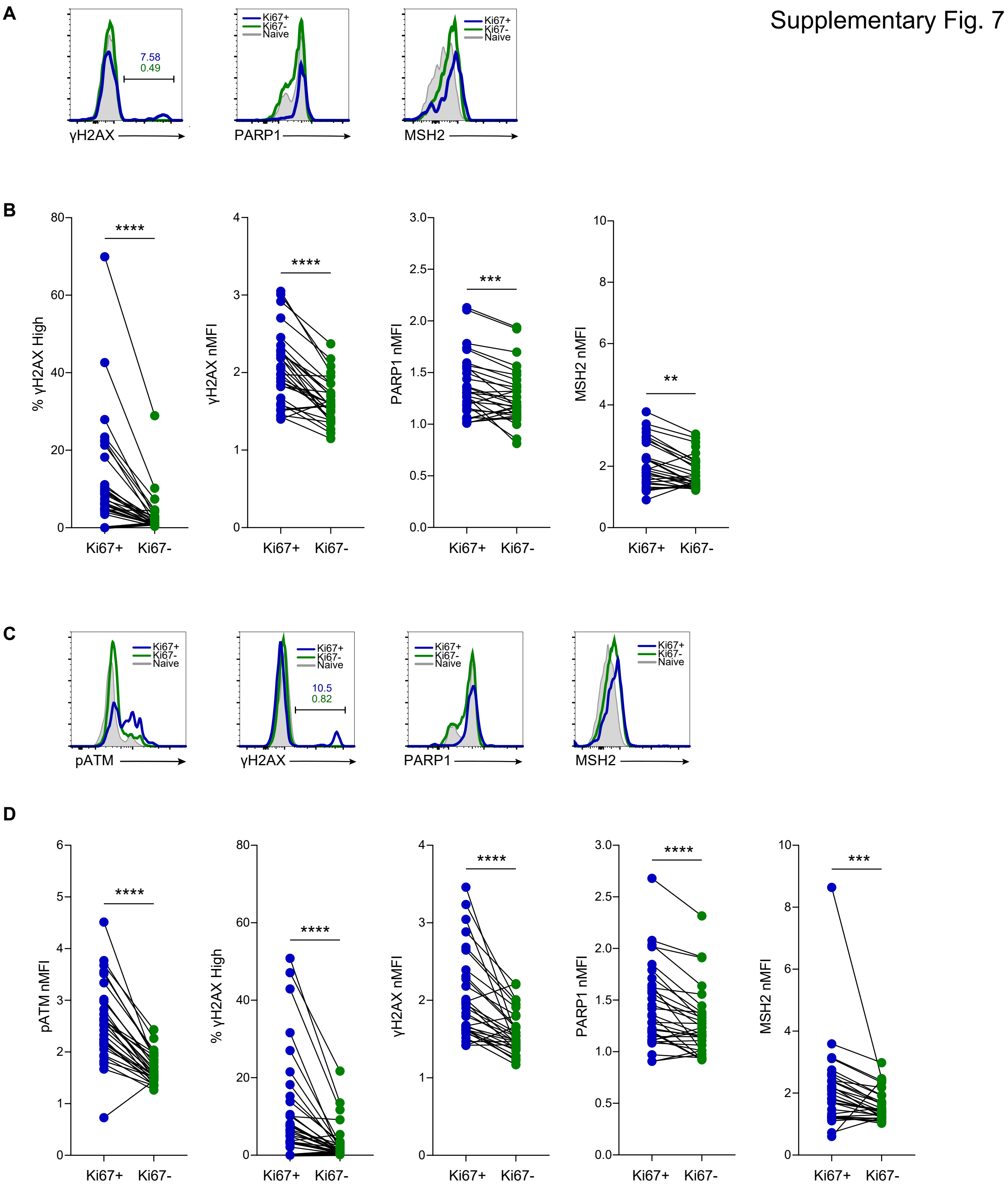
Supplementary Fig. 7
A
γH2AX
7.58
0.49
PARP1
MSH2
Ki67+
Ki67-
Naive
B
% γH2AX High
γH2AX nMFI
PARP1 nMFI
MSH2 nMFI
****
***
**
Ki67+
Ki67-
C
pATM
γH2AX
10.5
0.82
PARP1
MSH2
D
pATM nMFI
% γH2AX High
γH2AX nMFI
PARP1 nMFI
MSH2 nMFI

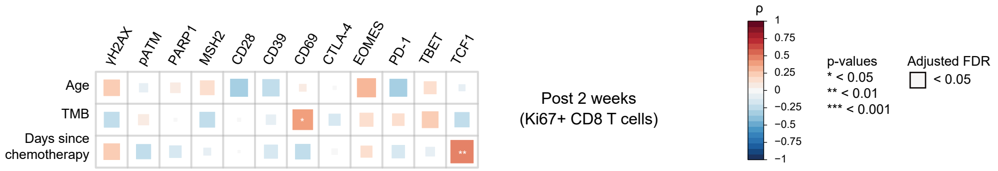
γH2AX
pATM
PARP1
MSH2
CD28
CD39
CD69
CTLA-4
EOMES
PD-1
TBET
TCF1
Age
TMB
Days since chemotherapy
*
**
Post 2 weeks
(Ki67+ CD8 T cells)
ρ
1
0.75
0.5
0.25
0
−0.25
−0.5
−0.75
−1
p-values
* < 0.05
** < 0.01
*** < 0.001
Adjusted FDR
< 0.05

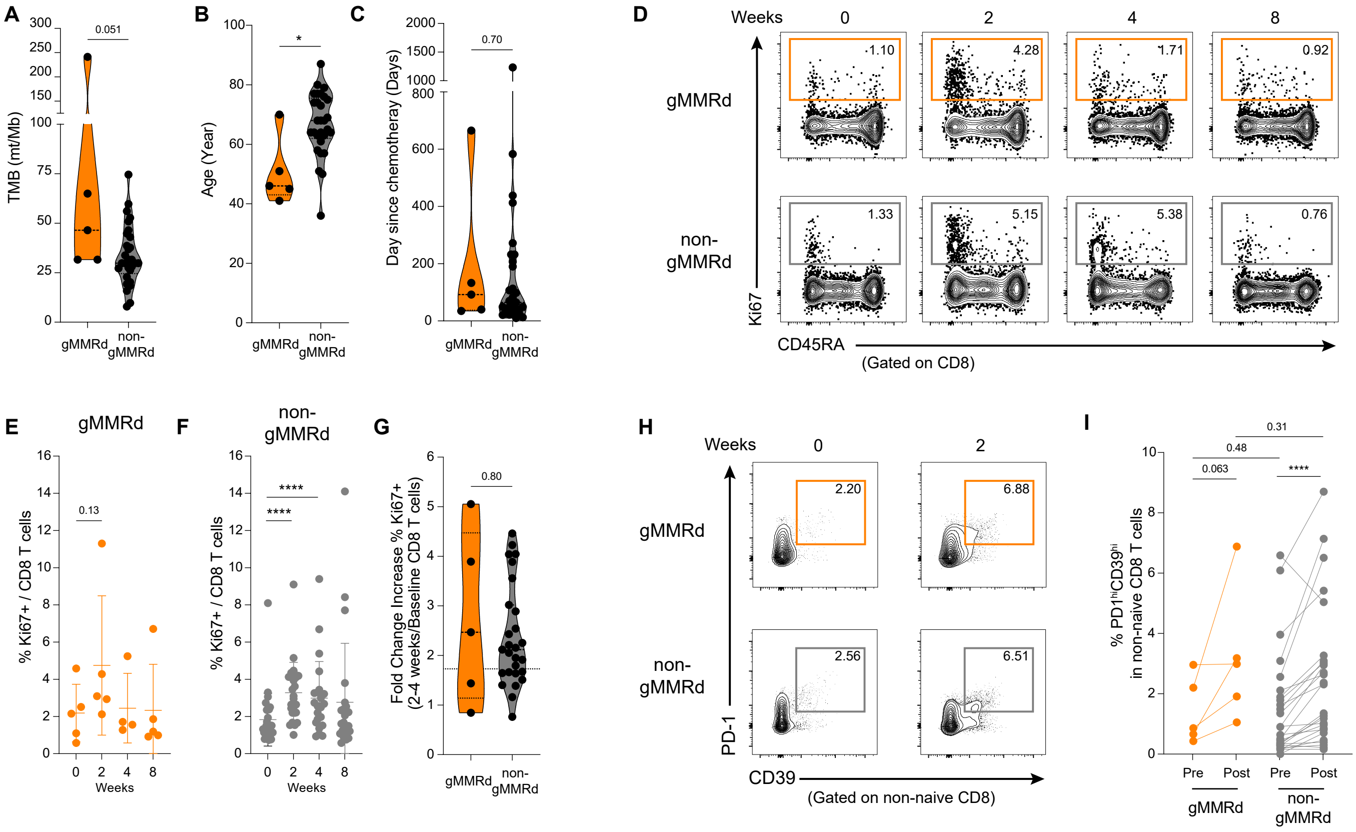
A
TMB (mt/Mb)
0.051
gMMRd
non-gMMRd
B
Age (Year)
*
C
Day since chemotheray (Days)
0.70
D
Weeks
0
2
4
8
gMMRd
non-gMMRd
1.10
4.28
1.71
0.92
1.33
5.15
5.38
0.76
Ki67
CD45RA
(Gated on CD8)
E
gMMRd
% Ki67+ / CD8 T cells
0.13
Weeks
F
non-gMMRd
****
G
Fold Change Increase % Ki67+
(2-4 weeks/Baseline CD8 T cells)
0.80
H
2.20
6.88
2.56
6.51
PD-1
CD39
(Gated on non-naive CD8)
I
% PD1hiCD39hi
in non-naive CD8 T cells
0.31
0.48
0.063
Pre
Post

Supplementary Fig. 10

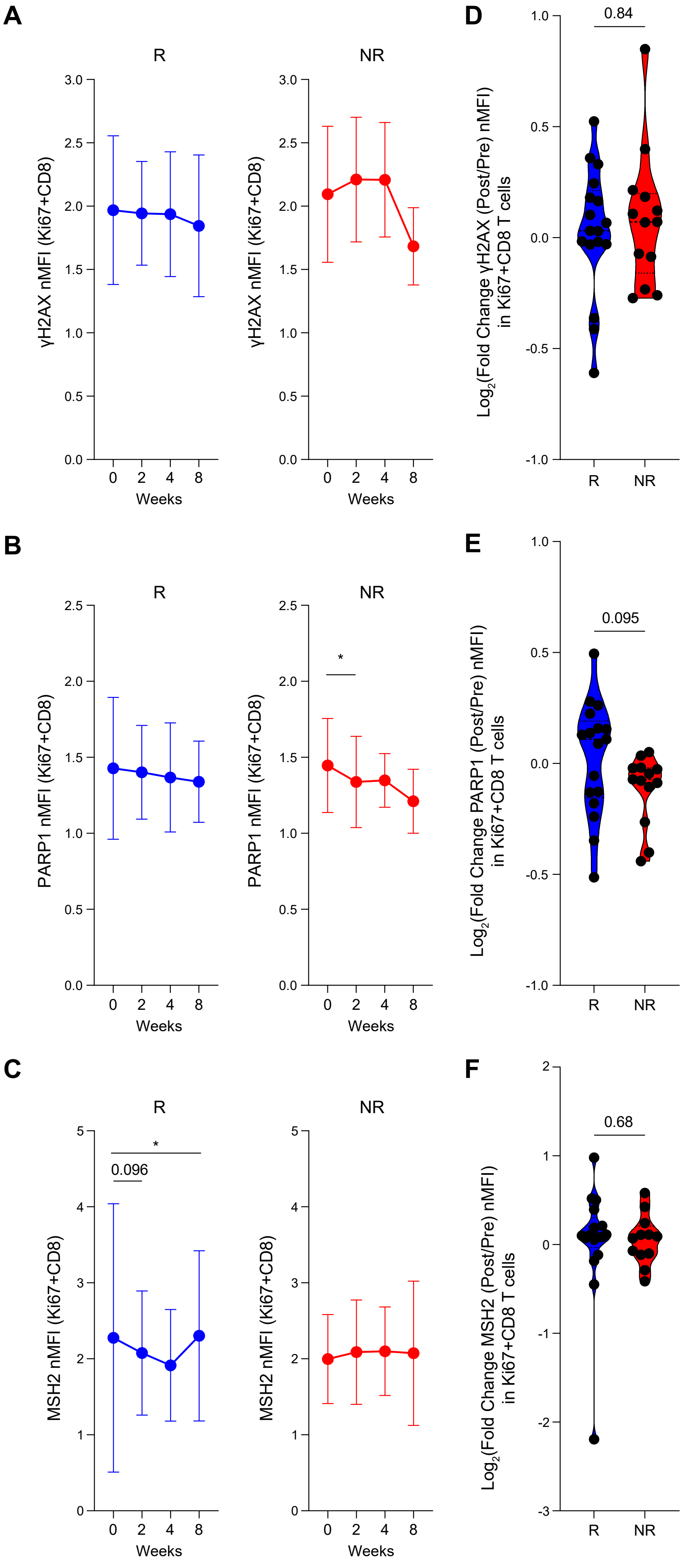

Supplementary Fig. 11

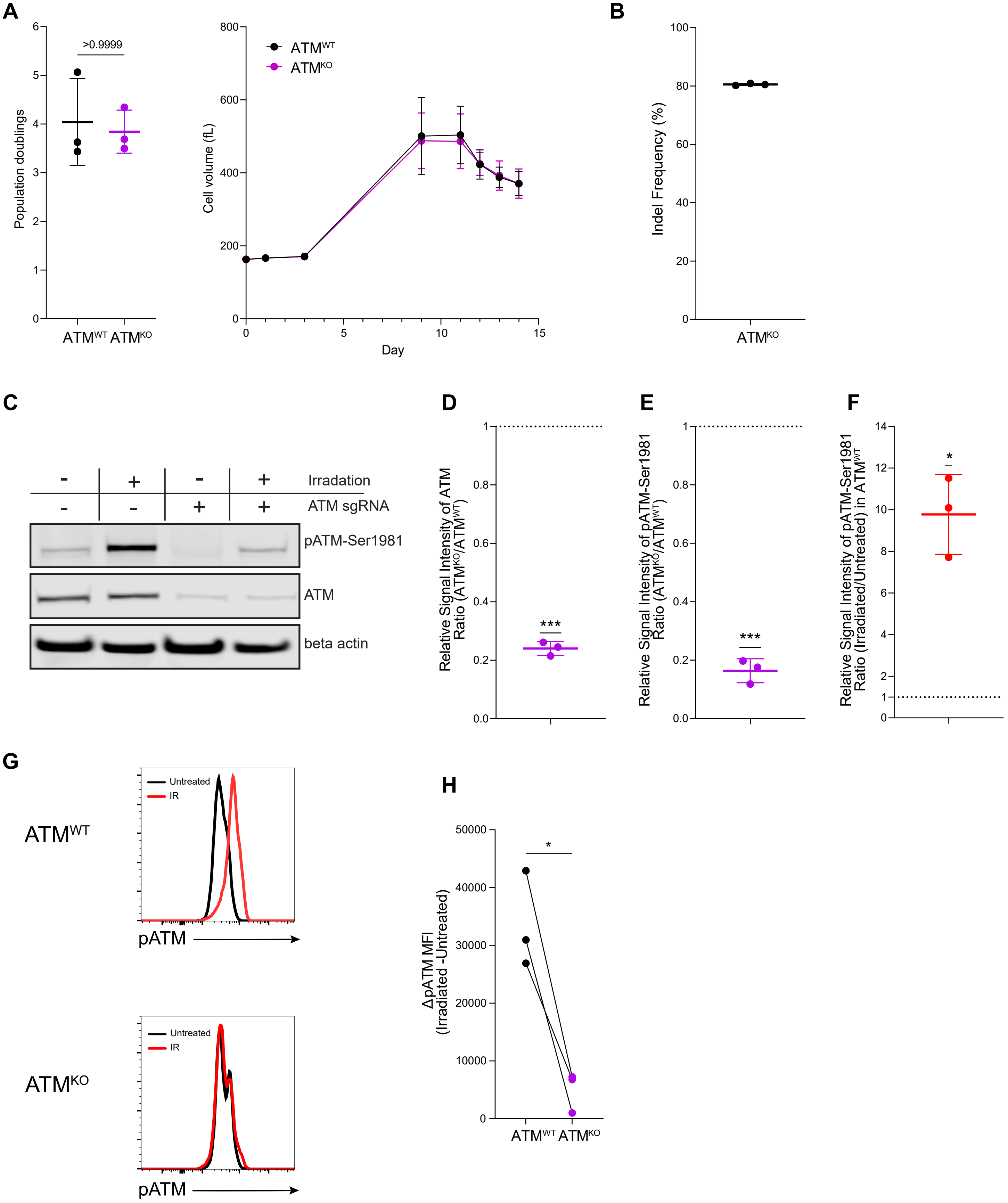

Supplementary Fig.12

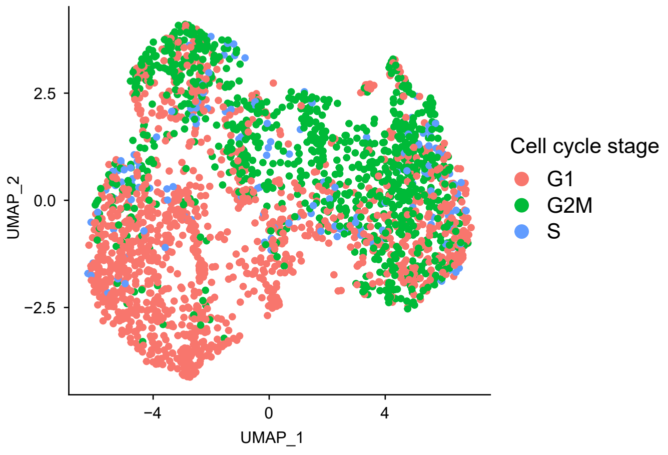

Supplementary Fig.13

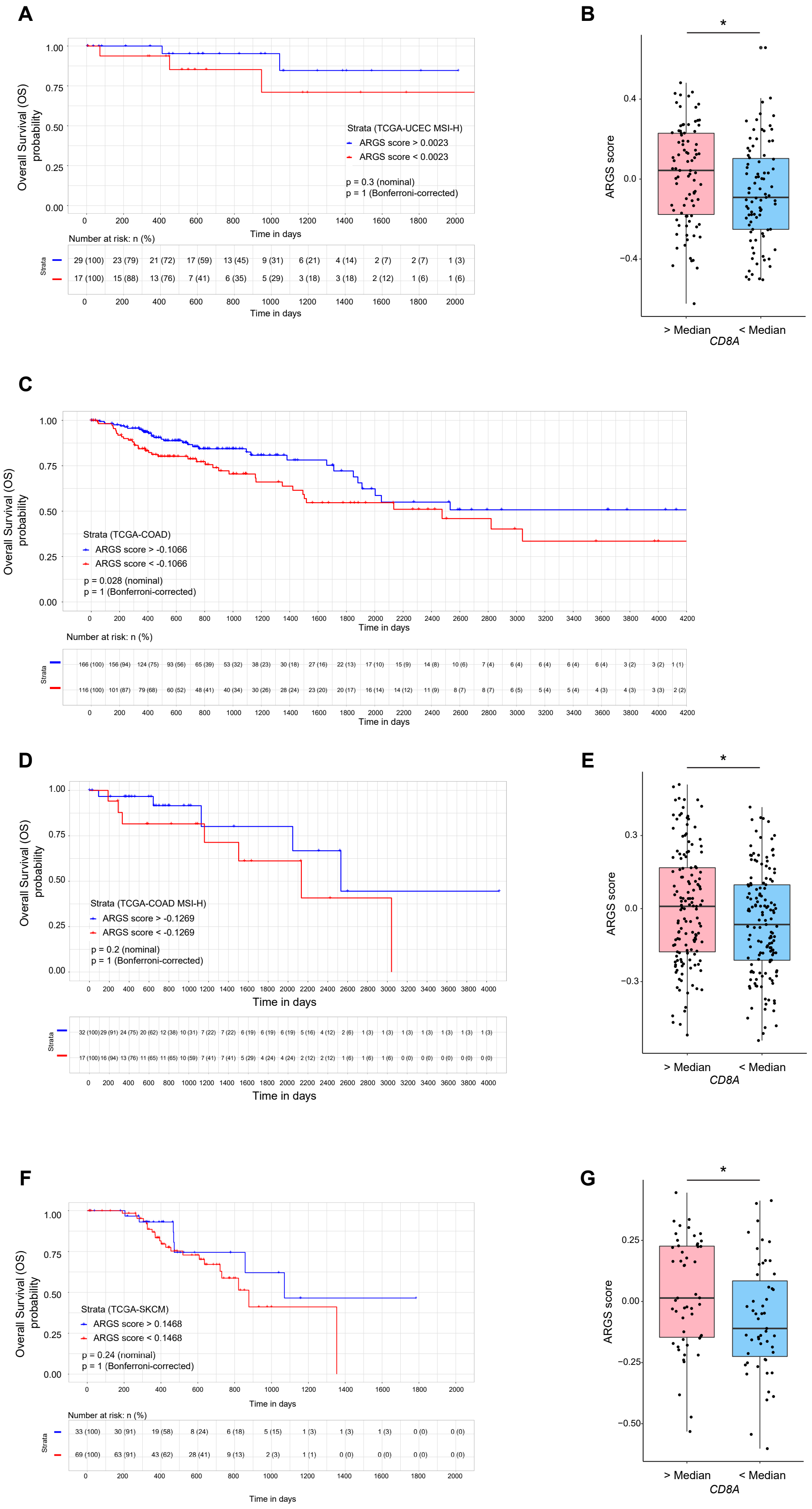
